# Prom1 and Notch regulate ciliary length and dynamics in multiciliated cells of the airway epithelium

**DOI:** 10.1101/2022.01.13.475833

**Authors:** Carlos F. H. Serra, Helu Liu, Jun Qian, Munemasa Mori, Jining Lu, Wellington V. Cardoso

## Abstract

Differences in ciliary morphology and dynamics among multiciliated cells of the respiratory tract have been well reported and known to contribute to efficient mucociliary clearance. Nevertheless, little is known about how phenotypic differences among multiciliated cells are established in the mammalian lung. Here we show that Prominin-1 (Prom1), a transmembrane protein widely used as stem cell and tumor-initiating marker, is crucial to this process. During airway differentiation, Prom1 becomes restricted to multiciliated cells, where it is expressed at distinct levels along the proximal-distal axis of the airways and in the adult airway epithelium in vitro. We found that Prom1 is induced by Notch in post-specified multiciliated cells and that Notch inactivation abolishes the gradients of Prom1 in the developing airways and in differentiating organotypic cultures. Prom1 was not required for multicilia formation and when inactivated resulted in longer cilia, which remained functional but beating at a lower frequency. Disruption of Notch resulted in opposite effects and suggested that Notch fine-tunes Prom1 levels to regulate the multiciliated cell phenotype and generate diversity among these cells in the respiratory tract. By controlling these features, this mechanism contributes to the innate defense of the lung against environmental agents and prevent pulmonary disease.

**Significance Statement:** Multiciliated cells are integral components of the epithelia from a variety of organs. In the respiratory tract they are crucial for mucociliary clearance, a first line of defense against environmental agents and microorganisms. Regional differences in ciliary morphology and dynamics of multiciliated cells have been well described. However, little is known about the events generating phenotypical and functional differences among these cells in airways. Here we provide evidence of a novel mechanism in post-specified multiciliated progenitors whereby local Notch and Prom1 regulate ciliary length and ciliary beating to generate morphological and functional diversity among the multiciliated cells. The findings provide insights into the impact of these signals in maintaining the integrity and function of the airway epithelium, preventing pulmonary disease.

## Introduction

Cilia are highly conserved organelles formed by microtubule-based apical protrusions, the axonemes, that anchor to the cytoplasm via the basal bodies (1-3). Nearly every cell has a single non-motile cilium, the primary cilium, critical for transducing cell-sensing signals (4). By contrast, motile cilia have a more restricted tissue distribution and are usually found in epithelia that specialize in fluid movement, such as in the neural tube, the fallopian tubes, and the conducting airways of the respiratory system (5). In these tissues, motile cilia are present in large numbers at the surface of multiciliated cells (100-200 per cell), often found interspersed with secretory cells (6, 7).

In the respiratory system, multiciliated cells are present throughout the entire air-conducting passages, from the trachea to the terminal bronchioles, where they are responsible for mucociliary clearance (8). The synchronized beating of cilia across multiciliated cells continuously propels upwards the mucous layer produced by the secretory cells, thus protecting the lungs against inhaled pollutants and pathogens. Along with the physical barrier formed by the tight junctions between epithelial cells, these mechanisms are essential components of the innate immunity (9). Disruption of these mechanisms by ciliary defects, such as ciliopathies, or by abnormal mucous production is frequently associated with respiratory infections and chronic obstructive pulmonary disease (10, 11).

Several studies have described regional differences in the phenotype and ciliary dynamics of multiciliated cells along the respiratory tract of the mammalian lung. Analysis of mucociliary clearance has shown a progressive decrease in the velocity of mucous transport across different generations of conducting airway from the trachea to bronchioles (12). Multiciliated cells also display marked regional differences in ciliary length with longer cilia reported predominantly in the trachea and extrapulmonary bronchi when compared to those in the terminal bronchioles (6). These differences are already evident during late gestation, and suggest the influence of distinct local contexts during multiciliated cell differentiation (8). Studies in primary cilia have implicated a large number of pathways in the regulation of cilia assembly and growth (13); however, less is known about the mechanisms generating phenotypical differences in the multiciliated cells of the airways.

There is evidence that the Prominins, a family of transmembrane glycoproteins that localize to plasmalemmal protrusions, including that of microvilli and cilia, influence ciliary morphology and function in different biological systems (14). Prominin-1 (Prom1, CD133) has been widely used as a stem cell marker in various tissues, and is considered to be a key marker of tumor-initiating cells in cancers. PROM1 is expressed in the adult human lung, with high levels reported in non-small cell carcinomas associated with poor prognosis (15). It is of particular interest that, in the adult lung, Prom1 is also found to be paradoxically expressed in the multiciliated cells of the airways, a cell type known to be terminally differentiated (16-18). The association of Prominins with cilia appears to be well-conserved, as loss-of-function of these proteins results in various cilia-related phenotypes, including retinal degeneration and left-right asymmetry in non-mammalian species (14, 19). Interestingly, some of the Prom1-related ciliary phenotypes have also been linked to defects in Notch-signaling, a pathway involved in the development of both monociliated and multiciliated cells (14, 20, 21). In fact, Prom1 and Notch are known to interact in different cell types and contexts to regulate cellular behaviors (22). These observations raise intriguing questions about the functional significance of Prom1 in multiciliated cells, the processes mediated by Prom1 and Notch-signaling, and their potential involvement in the generation of multiciliated cell diversity in the mammalian respiratory tract.

Here, we provide evidence that, during differentiation of the airway epithelium, Prom1 becomes restricted to multiciliated cells, where it is crucial in controlling cilia length and ciliary dynamics. Using mouse genetic approaches and functional assays in organotypic cultures we show that Prom1 and Notch-signaling regulate these features, ultimately establishing differences in morphology and function among multiciliated cells. Furthermore, we show that, by acting in multiciliated cells, Prom1 levels are key in the modulation of mucociliary clearance. Our results identify Prom1 as an integral regulator of the multiciliated cell function likely to contribute significantly to the innate defense mechanisms of the lung.

## Results

### Prom1 becomes restricted to multiciliated cells in differentiating airway progenitors

Prom1 has been widely described as a stem cell marker, but has also been shown to be expressed in the differentiated epithelia of a variety of organs (23-25). In a previous transcriptome analysis we found Prom1 expression enriched in a population of non-secretory cells of late gestation lungs (below, 29). Despite having been reported in the developing lung (26), information about the Prom1 spatial and temporal patterns of expression was unavailable. We identified Prom1 expression uniformly distributed throughout the epithelium of the embryonic E14.5 respiratory tract, from the trachea to the distal buds (Fig. 1A, Fig. S1A). At this stage, the airway epithelium is still largely undifferentiated, thus suggesting that *Prom1* marked these cells at their progenitor stem-like state. *Prom1* transcripts were also detected in the epithelia of other developing tissues, including the kidney, brain, and intestine (Fig. S1A). Immunofluorescence assays (IF) of the E14.5 airway epithelium further revealed the Prom1 subcellular localization restricted to the apical plasma membrane domain of the epithelial tubules (Fig. 1A). As the airway epithelium underwent differentiation, Prom1 was silenced in secretory cells, while remaining strongly expressed in multiciliated cells. IF of E18.5 lungs showed Prom1 double-labeling together with Foxj1, but not with CC10 (markers of multiciliated and secretory cells, respectively) (Fig.1B, Fig. 1C). Intriguingly, at this stage, Prom1 labeling was not only restricted to multiciliated cells, but its signal intensity was regionally distinct throughout the respiratory tract epithelium. Plotting of the fluorescence intensity from different airway generations of E18.5 lungs showed a gradient of Prom1 expression along the proximal-distal axis of the airways, with lower levels in the trachea and increasing signal intensity in intrapulmonary airways (Fig.1D).

**Figure 1.**
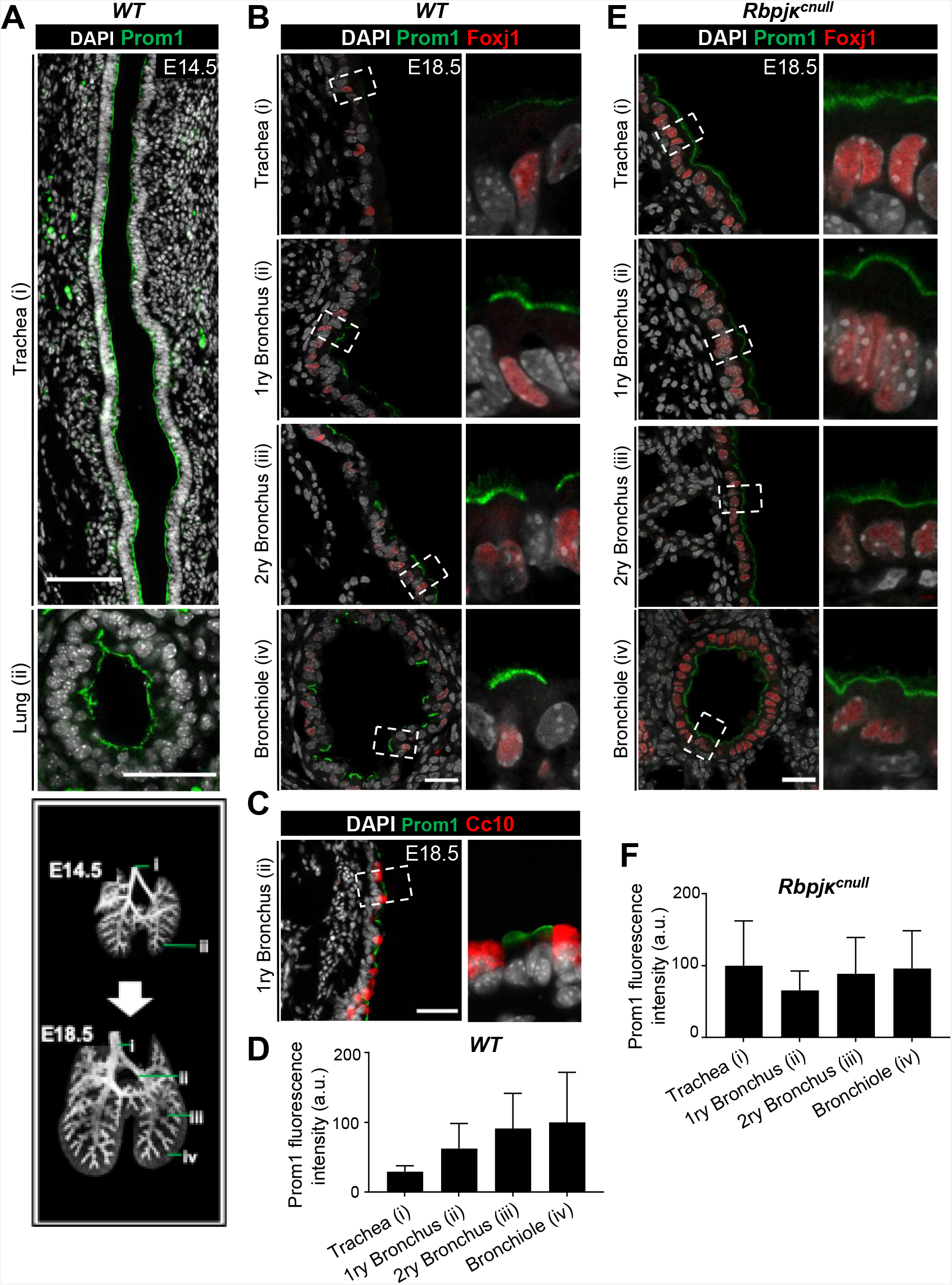
Prom1 becomes restricted to multiciliated cells and its regional levels are controlled by Notch signaling in the developing lung. **(A)** Immunofluorescence (IF) of Prom1 in the E14.5 trachea and intrapulmonary airways at the sites indicated by the numbered green bars in the diagram at the bottom left (adapted from Alanis et al. (49), depicting the stages analyzed in panels A-C, E). Prom1 is detected throughout the epithelium at the apical surface. **(B)** Double Prom1/Foxj1 IF in E18.5 wild type lungs. Each panel represents a series of single optical sections from different airway generations as depicted in the diagram (Boxed areas enlarged on the right panels). Prom1 is detected in multiciliated cells marked by Foxj1; labeling is stronger in intrapulmonary airways. **(C)** Double IF of Prom1/Cc10 in E18.5 wild type lungs. Boxed areas are enlarged on the right panel. Prom1 is not detected in secretory cells (marked by Cc10). **(D)** Graph: Prom1 fluorescent labelling intensity in different airway generations of E18.5 wild type lungs. Bars represent the mean + SD (arbitrary units, values normalized as percent mean fluorescence intensity in bronchioles). Prom1 proximal-distal gradient with higher expression in bronchioles. **(E)** Double Prom1/Foxj1 IF in E18.5 Rbpjκ^cnull^ lungs in different airway generations as depicted in diagram (green bars, bottom left). Boxed areas are enlarged on the right panels. Airways are overpopulated by multiciliated cells uniformly labelled by Prom1. **(F)** Graph: Prom1 fluorescent labelling intensity in different airway generations of E18.5 Rbpjκ^cnull^ lungs. Bars represent the mean + SD (values normalized as percent mean fluorescence intensity in trachea). DAPI is gray in all panels. 1ry = Primary; 2ry = Secondary. Scale bars in A (top and bottom panels), B, C and E = 100 μm and 50 μm, 25 μm, 50 μm and 25 μm, respectively.

We asked whether Prom1 could have a similar cell-type restriction during the differentiation of adult airway progenitors. Thus, we isolated airway basal cells from adult murine tracheas and expanded them to confluency in Transwell cell culture plates, as previously reported (27). IF-confocal microscopy showed Prom1 expressed uniformly in all basal cells, largely at the apical surface, a pattern reminiscent of that found in the progenitors of the developing E14.5 airway epithelium in vivo (Fig. S1B, Fig. 1A). Analysis of these cultures differentiating under air-liquid interface (ALI) conditions showed Prom1 selectively detected in multiciliated cells, as confirmed by the co-labeling with β-IV Tubulin (Fig. S1C). Notably, these cells displayed a great degree of heterogeneity in regards to Prom1 expression, as seen by the wide range of labeling intensities throughout culture period (Fig. S1C-D,). We reasoned this could be ascribed to potential differences in the stage of differentiation of the multiciliated cells arising in the same culture (timing), or to the presence of progenitor cells cultured from distinct regions of the trachea (spatial heterogeneity). The fact that the differences in Prom1 expression persisted in later-stage cultures when multiciliated cells were nearly all mature (ALI day 27, Fig. S1D suggested that the heterogeneity we observed reflected intrinsic differences in the developmental program of these progenitors associated with their original location. Overall, these results revealed a conserved pattern of restriction of Prom1 expression as cells transition from an undifferentiated stem-like status to a highly differentiated multiciliated cell lineage both in the developing lung and in the adult airways differentiating in vitro. They also underscore the substantial heterogeneity of multiciliated cells in regards to Prom1 expression, both *in vivo* and *in vitro*.

### Notch-signaling controls local and regionally distinct levels of Prom1 in multiciliated cells

To gain additional insights into the heterogeneity of multiciliated cells of the airway epithelium, we examined lungs from mouse mutants in which these cells had been massively expanded by disruption of Notch-signaling. Notch regulates cell fate decisions in the developing airways by fostering secretory cell fate and preventing excessive multiciliated cell differentiation (7). Analysis of E18.5 mutants in which we inactivated canonical Notch-signaling early in the lung epithelium (*ShhCre; Rbpjkf/f*, hereafter Rbpjk^cnull^) showed the expected expansion of multiciliated cells in the airways (Fig. S2)(7, 28)). Notably, Prom1/Foxj1 double-IF staining of E18.5 Rbpjk^cnull^ lungs not only showed airways massively overpopulated by double-labeled multiciliated cells, but also revealed an overall low uniform pattern of Prom1 expression in multiciliated cells throughout extra- and intrapulmonary airways (Fig. 1, Fig. S2A). Assessment of Prom1 fluorescence intensity from different airway generations of E18.5 mutant lungs showed similar values along the proximal-distal axis, in contrast to the graded distribution seen in wild type (WT) lungs (Fig. 1D, F). Thus, disruption of Notch-signaling altered Prom1 expression within these cells, attenuating the normal gradient observed in WT lungs. Transcriptome profiling of E18.5 Rbpjk^cnull^ lungs showed that, when Prom1 is normalized by Foxj1 (both selectively expressed in multiciliated cells) Prom1 levels were consistently lower in mutants compared to controls (29). This contrasted with the behavior of other cilia-associated gene markers which, when similarly normalized by Foxj1, were upregulated in Rbpjk^cnull^ lungs (Fig. S2B). The observation suggested that in differentiated multiciliated cells Notch is a positive regulator of Prom1. Remarkably, similar analysis of mice in which Notch signaling was constitutively activated in the lung epithelium (ShhCre; NICD) showed increased Prom1 expression normalized by Foxj1 in E18.5 lungs mutants compared to controls (Fig. S2B). Thus, Notch appears to not only control the abundance of multiciliated cells in the airways but also differentially regulate the levels of Prom1 regionally in these cells.

To examine whether Notch also influenced the program of Prom1 expression in adult progenitors undergoing multiciliated cell differentiation, we isolated basal cells from adult WT murine tracheas (as above) and disrupted Notch-signaling pharmacologically with the gamma-secretase inhibitor DAPT. Efficient Notch inhibition was confirmed by the expansion of basal cells in ALI day0 and excessive multiciliated cell differentiation by ALI day8 (30). DAPT treatment resulted in Prom1 signals slighted decreased at day 0 compared to controls but by day 8 Prom1 expression sharply declined in the DAPT cultures (Fig. 2A).

**Figure 2.**
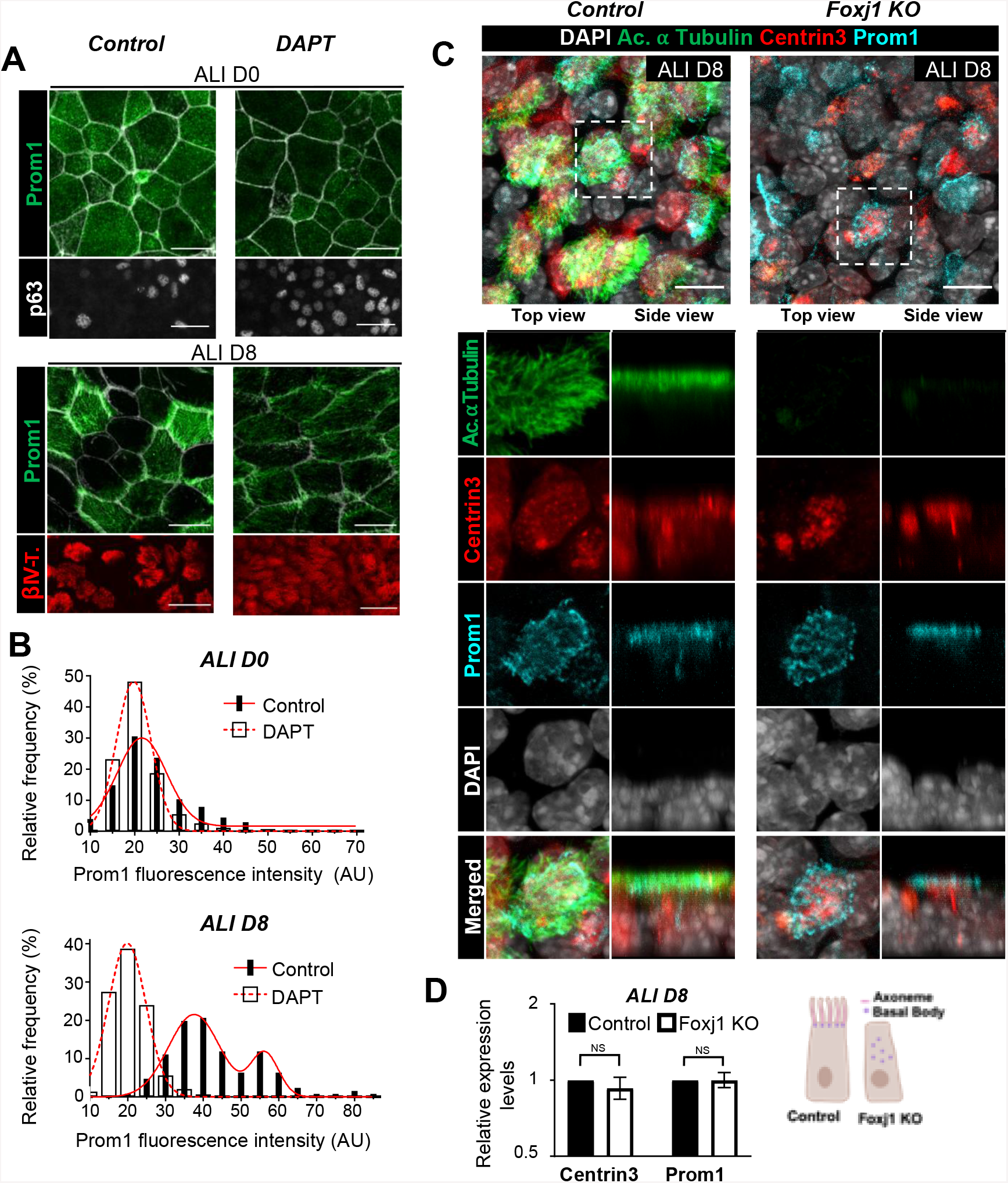
Prom1 expression is dependent on endogenous Notch signaling in multiciliated cells and independent of multicilia formation. **(A)** Immunofluorescence (IF) of Prom1, p63 and acetylated α-tubulin in control and DAPT-treated ALI airway cultures at days 0 and 8. Expansion of the basal (p63+) and multiciliated (acetylated α-tubulin+) cell population. Prom1 expression decreased by DAPT. Prominent heterogeneity of Prom1 signals in control ALI day 8 compared to day0 (Phalloidin shown in gray in top panels). **(B)** Bar charts representing the frequency distribution of the Prom1 labelling intensities in individual Prom1+ cells from maximum projections confocal micrographs of ALI cultures at days 0 and 8. The red lines represent non-linear curve fittings for the frequency distribution of control (DMSO, solid line) and Notch-deficient (DAPT, dashed line) ALI cultures. At day 0 the solid red line (Mean ± SD of 21.73±6.19 %, R^2^=0.9439) and the dashed red line (Mean ± SD of 19.76±3.90 %, R^2^=0.9882) represent single gaussian curves. At day 8 the solid red line represents the sum of two gaussian curves (Mean ± SD of 37.59±6.80 % and 56.24±3.68 %, R^2^=0.9874), and the dashed red line represents a single gaussian curve (Mean ± SD of 19.86±4.97 %, R^2^=0.9865). **(C)** Triple IF of Prom1, acetylated α-tubulin and Centrin3 in ALI day 8 cultures incubated with CRISPR/Cas9 lentiviral vectors carrying a scrambled-sequence (Control, left panels) or a Foxj1-targeting sequence (Foxj1 KO, right panels). Top and side views of the boxed areas are enlarged in bottom panels for each marker. All panels represent maximum intensity projections. In Foxj1 KO cultures the ciliated cell fate is specified but multiciliogenesis does not occur; Prom1 expression is unaffected and still limited to the apical region of multiciliated-fated cells (marked by Centrin3). DAPI is gray in all panels. **(D)** qPCR analysis of Prom1 and Centrin3 in Control and Foxj1 KO day 8 ALI cultures (N=3). There is no significant difference in Prom1 and Centrin3 in Foxj1 KO. Graph represents fold change ± CI95 on a log2 scale. Student’s t-test: ns, not significant. Scale bars in A and D = 25 μm and 10 μm, respectively.

As noted before, in control day 8 cultures Prom1 was expressed in a much wider range of fluorescence intensity among the multiciliated cells than in the control basal cells at day 0 (Fig. S1B-C, Fig.2A left panels). Indeed, the frequency distribution of Prom1 signal intensity in day 0 control cultures could be represented by a single Gaussian curve. This contrasted with the pattern seen in the control day 8 cultures, which could be better explained by the sum of two Gaussian curves (R2=0.9874) rather than by a single one (R2=0.8326) (Fig. 2B). The pattern suggested the underlying presence of at least two distinct multiciliated populations based on differences in Prom1 expression level of control cultures. Interestingly, in DAPT-treated cultures at day 8 the frequency distribution of Prom1-fuorescence intensity in multiciliated cells was found in a single peak, in contrast to the double peak seen of respective controls (Fig. 2B). Thus, disruption of Notch not only decreased Prom1 levels but also led to the appearance of a more uniform population of Prom1^low^ cells. Taken together, these results show that multiciliated cells in the airway epithelium differ in their ability to express Prom1. This appears to depend on their distinct local or regional contexts in the respiratory tract *in vivo* and in vitro as we observed in E18.5 lungs and in adult organotypic cultures. These differences seem to also depend on Notch-signaling, suggesting that this pathway may be differentially activated in subpopulations of multiciliated cells.

### Prom1 is not required to initiate multiciliogenesis and is independent of multicilia formation to be expressed

Our data showed that, during airway epithelial differentiation, Prom1 expression is selectively maintained in the cells committed to initiate the program of multiciliogenesis. This led us to examine whether preventing subsequent events of the ciliogenesis program in cells fated to become multiciliated would have any impact in Prom1 expression and localization. Foxj1 is a transcription factor crucial for initiation of multiciliogenesis once large-scale centriole amplification has taken place, for it regulates the docking of hundreds of centrioles (basal bodies) to the apical surface of multiciliated cells, from which axonemes will elongate. Foxj1 loss-of-function results in failure to localize basal bodies apically and properly form axonemal ciliary structures at the cell surface (31). We employed a Crispr/Cas9 gene-editing approach to disrupt the expression of Foxj1 in airway progenitors *in vitro*. A lentiviral vector carrying an sgRNA targeting Foxj1. Cas9, and a puromycin resistance gene was used to create double-stranded breaks in the Foxj1 coding sequence of airway progenitors, ultimately resulting in the knock down of functional Foxj1 protein in ALI cultures (Fig. S3A). A lentiviral vector containing a scrambled sgRNA sequence was used as control.

Freshly-isolated adult airway progenitors were incubated with lentivirus-containing medium for 48 hours. Transduced cells were selected by puromycin resistance, expanded to confluency, and finally differentiated under ALI conditions for 8 days, as previously reported (32). Efficient knockdown of Foxj1 was demonstrated by qPCR and cytometry analysis, with both showing a significant decrease in Foxj1 expression, when compared to controls (Fig. S3B-C). Disruption of Foxj1 expression resulted in the expected loss of acetylated α-tubulin-labeled axonemes characteristic of multiciliated cells. Nonetheless, the multiciliated identity of these cells could still be readily recognized by the abundant expression of Centrin3, which labels the multiple basal bodies that remained undocked in the absence of functional Foxj1 (23) (Fig. 2C). Remarkably, IF of Foxj1-disrupted cultures showed Prom1-expression still limited to the Centrin3+ cells, which were committed to the multiciliated cell fate. Furthermore, Prom1 apical localization remained unaffected in spite of the defect in basal body docking and the consequent lack of cilia (Fig. 2C). Morphometric and qPCR analyses revealed no significant differences in the expression levels of *Prom1* and *Centrin3* in Foxj1-disrupted cultures, when compared to controls (Fig. 2D). Together, these results indicate that, although in the airway epithelium Prom1 is normally restricted to multiciliated cells, its expression and apical subcellular localization do not depend on basal body docking or elongation of axonemes.

### Disruption of Prom1 results in multiciliated cells with overly elongated cilia

Previous studies in *Prom1*^*−/−*^ mice have reported complex phenotypes, with varying severities depending on the genetic background. Some of the abnormalities include: retinal degeneration (33), spermatogenesis defects, abnormal intestinal-crypt proliferation and inflammation (34), and tooth developmental defects (35), among others. No information was available regarding the impact of Prom1 deletion in the airway epithelium. To gain insights specifically into this issue and circumvent the systemic effects of *Prom1*^*−/−*^ mice, we disrupted Prom1 expression in airway epithelial progenitors from adult mouse tracheas using a similar CRISPR/Cas9 approach (described above). Efficient disruption of Prom1 expression in airway progenitors was confirmed by qPCR, Western blot, and IF analyses (Fig. S4A-C). Immunostaining for p63, a transcription factor that marks the airway basal cells (36), indicated that the loss of Prom1 did not have any impact in the relative abundance of this population, nor in their ability to expand and form a confluent cell layer (Fig. S4D). In fact, confluence was reached just as fast in Prom1-CRISPR/Cas9 targeted cultures as it was in controls. Next, we examined the phenotype of multiciliated cells derived from the Prom1-deficient progenitors in ALI cultures. Loss of Prom1 had no effect in the balance of multiciliated vs secretory cells. We did not find significant differences in the relative expression levels of *Centrin3* and *Scgb3a2*, or in the number of Centrin3-labeled cells, when compared to controls (Fig. 3, Fig. S5). Analyses of fully differentiated cultures at a later stage (ALI day 27) showed a marked increase in cilia length in multiciliated cells from Prom1-deficient cultures compared to controls. XZ optical-reconstructions from confocal laser scanning micrographs identified long, Prom1-negative, acetylated α-tubulin-labeled axonemal structures (Fig. 3A). Morphometric analysis of cilia from scraped multiciliated cells, using an established method based on High-Speed Video Microscopy (HSVM) recordings of live cells (see Methods) (37), confirmed the increase in cilia length (CL) of multiciliated cells from Prom1-deficient cultures, when compared to controls (Prom1 KO Mean±SD = 111.2±3.6 %; *p* value = 0.0279; n = 5 individual cilia per multiciliated cell) (Fig. 3B).

**Figure 3.**
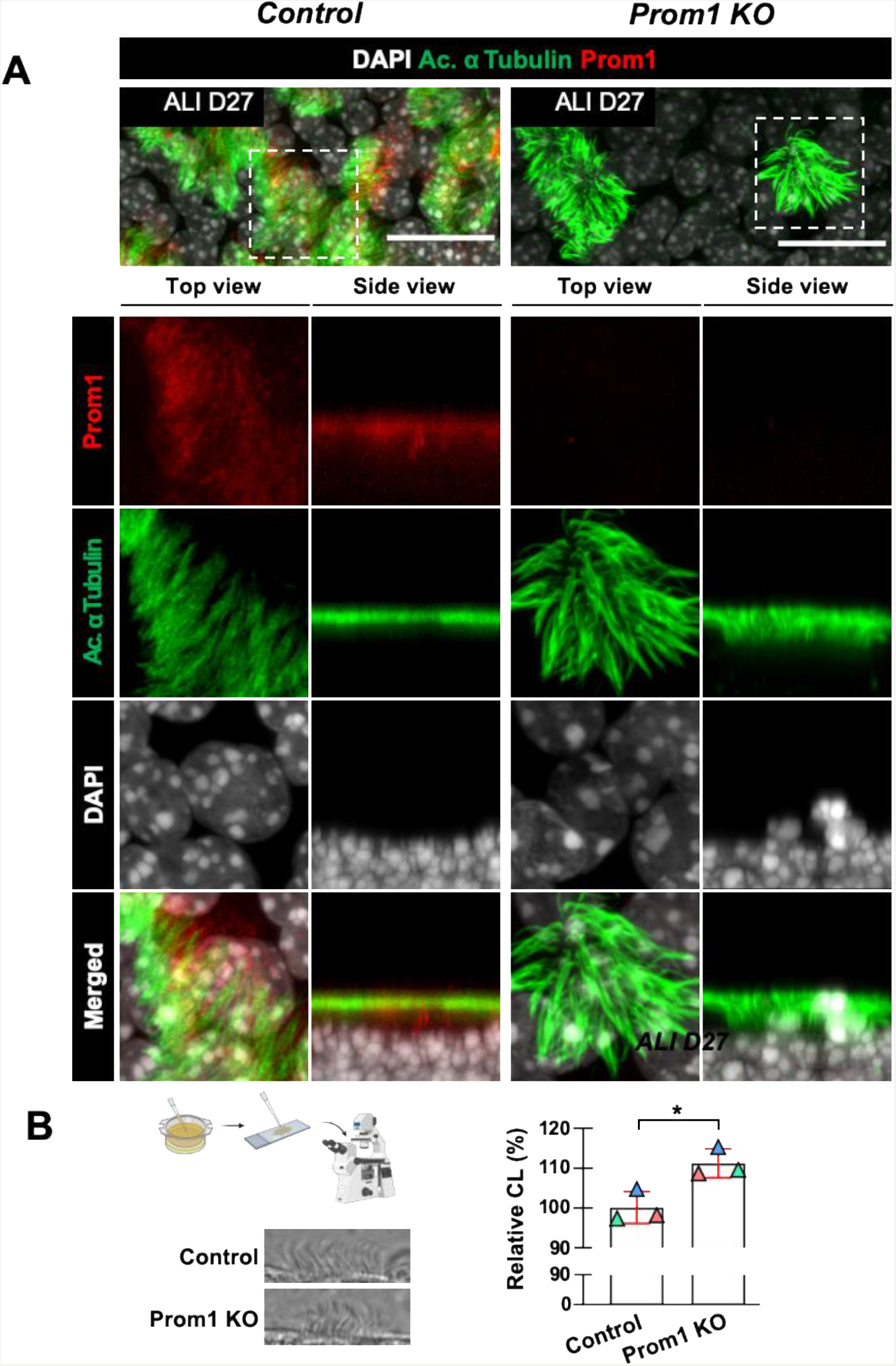
Disruption of Prom1 increases ciliary length (CL) in multiciliated cells. **(A)** Double immunofluorescence of Prom1 and acetylated α-tubulin in ALI day 27 cultures incubated with CRISPR/Cas9 lentiviral vectors carrying a control (left panels) or Prom1-targeting sequence (Prom1 KO, right panels). Top and side views of the boxed areas are enlarged in bottom panels. All panels represent maximum intensity projections. Prom1 KO does not prevent multiciliated cell differentiation but results in increased cilia length. **(B)** Cilia Length analysis from multiciliated cells of control and Prom1 KO ALI day 27 cultures. Top left: diagram methodology: airway epithelial cells scraped from Transwell membranes placed on a slide for imaging with a high-speed camera, and subsequent tracing of cilia. Bottom left: single, representative video frames for the conditions tested. Right: beeswarm plot of the well-means of relative cilia length (values normalized as percent mean cilia length of controls) superimposed on a bar chart representing the sample-mean ± SD. The original values were measured, in pixels, by averaging the length of 5 individual cilia from each multiciliated cell. Cilia length is increased in multiciliated cells from Prom1 KO cultures. Student’s t-test: *, *p* value < 0.05. DAPI is gray in all panels. Scale bars in A = 20 μm. Illustration created with BioRender.com.

### Ciliary beating and mucociliary transport are impaired in Prom1-deficient airways

Our analysis showed that the disruption of Prom1 expression in airway progenitors had no impact in their ability to undergo multiciliogenesis. However, it was unclear whether the abnormally long cilia found in Prom1-disrupted ALI were still functional and able to beat at frequencies comparable to those found in control cultures. To investigate this, we recorded HSVM videos of live, fully differentiated (ALI day 27) airway epithelial cultures that had been transduced with either a control, or a Prom1-disrupting lentiviral vector. The cells were imaged *en face* while still attached to the membranes, in 25 random fields per well, at a frame rate of 360 frames per second and at stable temperature of 37°C, as described by Oltean *et al* (37). The fast Fourier Transform (FFT) was used to calculate the Ciliary Beating Frequency (CBF) from the movement detected for each individual pixel of a video frame. Prom1-deficient multiciliated cells showed significantly lower CBF compared to controls (Prom1 KO: 37.52±2.17 Hz *vs*. Control: 42.46±1.78 Hz, adjusted *p* value = 0.0457). Thus, disruption of Prom1 in airway progenitors resulted in multiciliated cells with increased cilia length and lower frequencies of ciliary beating (Fig. 4A).

**Figure 4.**
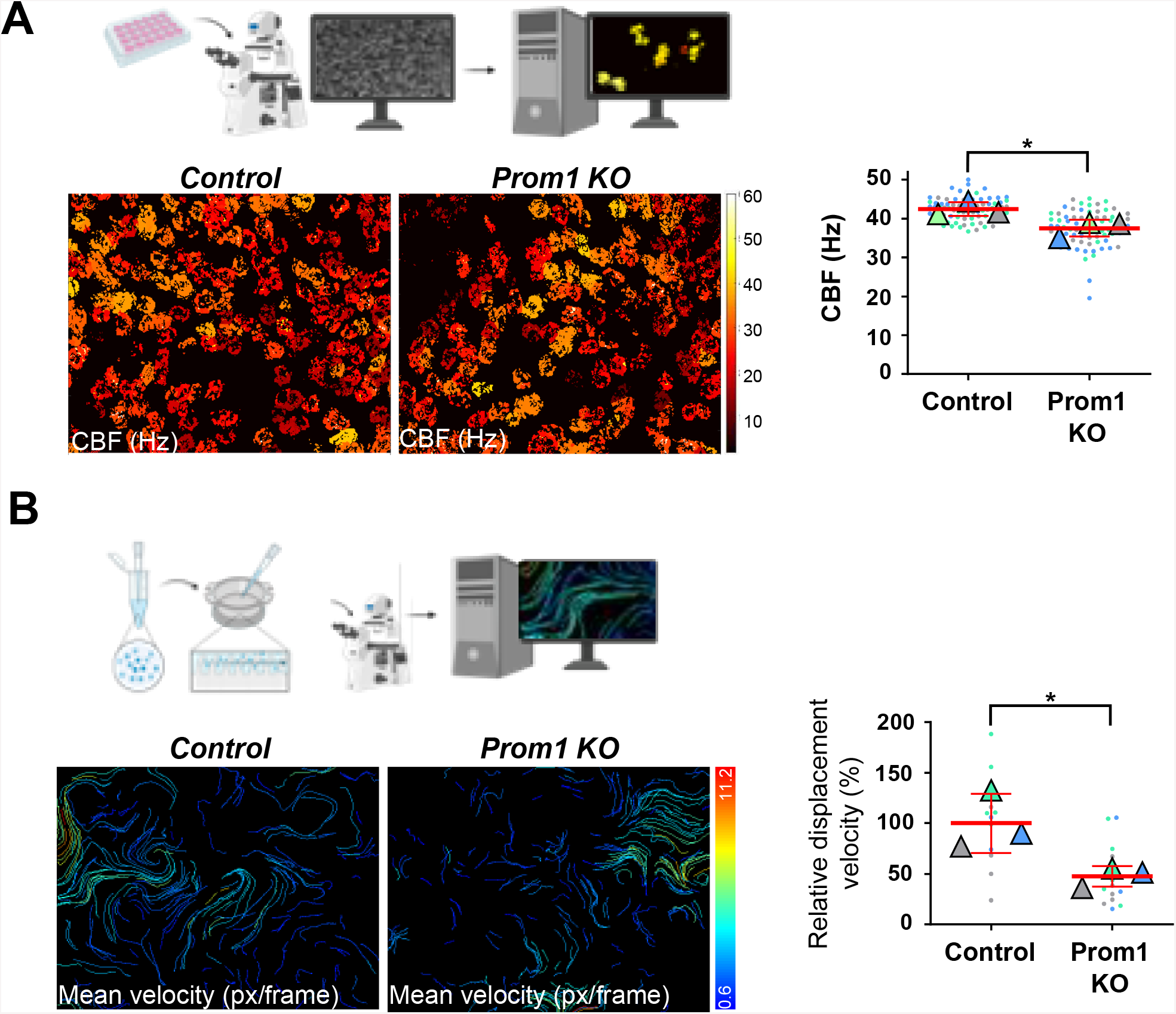
Loss of Prom1 impairs ciliary beating and mucociliary transport. **(A)** Analysis of Ciliary Beating Frequency (CBF) in ALI day 27 cultures incubated with CRISPR/Cas9 lentiviral vectors carrying control or Prom1 KO sequences. Top diagram depicts the methodology: live airway epithelial cells imaged in Transwell dishes with a high-speed camera and beating cilia analyzed using fast Fourier Transform. The heatmaps in the bottom left depict the CBF values of each pixel from representative recordings of Control and Prom1KO cultures. Right panel: beeswarm plot of the average CBF (Hz), of each video (●, N=25) and well (Δ, N=3), with the sample-means ± SD superimposed in red. Prom1 KO cultures had a significantly lower CBF compared to control. Student’s t-test: *, *p* value < 0.05. **(B)** Mucociliary Transport (MCT) in ALI day 27 cultures incubated with CRISPR/Cas9 Control or Prom1 KO vectors. Top diagram depicts methodology: fluorescent microspheres were added to the surface of live ALI cultures and imaged with a high-speed camera, the movement of the microspheres in the recordings was analyzed with TrackMate. The heatmaps in the bottom left depict the mean displacement velocity values of each fluorescent microsphere’s track, in pixel/frame, from representative recordings of Control and Prom1KO cultures. Right panel: beeswarm plot of the relative mean-displacement velocity weighted according to displacement distance of each respective track, of each video (●, N=5) and well (Δ, N=3), normalized so that the mean velocity of control = 100%, and with the sample-mean ± SD superimposed in red. Microspheres from Prom1 KO cultures had a significantly lower displacement velocity when compared to control. Student’s t-test: *, *p* value < 0.05. Illustration created with BioRender.com.

Since both ciliary morphology and dynamics differ along generations of airways *in vivo* and both were significantly altered in Prom1 KO cultures, we examined whether under control conditions, differences in CL among multiciliated cells are necessarily accompanied by corresponding changes in CBF. CBF and CL were measured together in HSVM recordings of multiciliated cells from control ALI day 27 cultures. Pearson’s correlation analysis showed no significant correlation between CL and CBF ([*r*] = -0.3521, *p* = 0.0994) (Fig. S6), in agreement with similar observations in ALI organotypic cultures from human donors (37). This suggested that the changes in ciliary length and dynamics mediated by Prom1 are unlikely to be interdependent and possibly results from Prom1 acting in multiciliated cells through different mechanisms.

Our analysis of airway organotypic ALI cultures showed Prom1 selectively expressed in multiciliated cells already early during differentiation. However, it was unclear when Prom1 started to influence CBF in these cells. Thus, we performed HSVM recordings in Prom1 KO cultures at different stages to investigate when Prom1 deficiency first altered ciliary dynamics. We found no difference in CBF in Prom1-deficient ALI cultures at day 8 compared to controls. However, two days later CBF was first significantly decreased in Prom1 KO cultures. Interestingly, similar analysis of CBF in ALI cultures treated with DAPT showed CBF significantly increased compared to controls already at day 8, suggesting that Notch acts on CBF even before Prom1, presumably through a Prom1-independent mechanism (Fig. S7).

To better understand the functional implications of Prom1-disruption in the airway epithelium, we assessed Mucociliary Transport (MCT) by tracking the displacement of fluorescent microspheres added to the surface of ALI cultures, as described by Oltean *et al* (37). Briefly, fluorescent microspheres were added to the surface of ALI day 27 cultures and HSVM recordings were performed which allowed us to track the displacement of individual microspheres, and subsequently determine the displacement distance (in pixels) and displacement velocity (in pixels/frame) using the TrackMate software (38). The final MCT value for each video was calculated by weight-averaging the displacement velocity of every track by their respective displacement distance. This analysis revealed a significant impairment of MCT in Prom1-deficient cultures (Prom1 KO Mean±SD = 47.61±10.09 %, *p* value = 0.0423) (Fig. 4B). Together, these results indicate that Prom1 influences various aspects of the multiciliated cell phenotype and behavior, which ultimately contribute to the efficiency of MCT. These observations place Prom1 as a relevant regulator of the multiciliated cell phenotype, and one who is likely to have a significant impact in the defense mechanisms of the airway epithelium.

## Discussion

Here, we provide evidence of Prom1 acting as a key regulator of the ciliary structure and function in multiciliated cells of the murine respiratory tract. We identified Prom1 widely distributed in the undifferentiated airway progenitors of both the developing and adult lungs, becoming restricted to multiciliated cells as differentiation takes place. Our findings of distinct levels of Prom1 in multiciliated cells raised the possibility that diversity of these cells is in part mediated by differential expression of Prom1. Our loss-of-function analyses showed that while Prom1 is not required for initiation of multiciliogenesis, it regulates ciliary length. We found that Prom1 operates together with Notch-signaling in the generation of multiciliated cell diversity in the developing and adult airways. Furthermore, we show that Prom1 contributes to generate functional differences among multiciliated cells by regulating ciliary dynamics, in a process independent of cilia length.

The precise mechanism by which Prom1 controls ciliary length in the airway epithelium is still unclear. Interestingly, analyses of mouse embryos during neurogenesis identified Prom1 in the plasmalemmal protrusions of primary cilia from neuroepithelial cells. The presence in the neural tube fluid of Prom1-rich membrane particles containing alpha tubulin suggests that cilia length could be controlled by shedding of Prom1-rich membrane domains (39). This mechanism is likely to be relevant in multiple biological contexts, since Prom1-containing membrane particles have been identified in a variety of body fluids, including urine, saliva, and seminal fluids (40). Studies in Madin-Darby Canine Kidney (MDCK) cells in which specific domains of Prom1 had been mutated suggest that the effect of Prom1 in cilia length can also involve interactions with ADP-ribosylation factor-like protein 13B (Arl13b), as well as cross-talking with cytoskeleton regulators in the cellular membrane microenvironment (14). Future work will clarify whether similar mechanisms are in place in airway multiciliated cells.

In our study, the functional relationship between Notch-signaling and Prom1-expression was intriguing. Although we had evidence of Notch regulation of Prom1, our data also suggested Notch-independent effects in multiciliated cells. We propose that, once airway progenitors have been committed to the multiciliated cell fate and ciliogenesis has initiated, subtle levels of Notch-activation in multiciliated cells regulate the expression of cilia-related genes, such as Prom1 to control ciliary morphology and function. This is supported by various lines of evidence: Notch-signaling is active and required to maintain proper ciliary length of the motile cilia in the zebrafish node, with Notch-inactivation resulting in shortened cilia and severe laterality defects in the embryo (20); Notch-activity in the murine developing neural tube has been shown to produce longer primary cilia (41). We have evidence that this is also true in differentiating multiciliated cells as seen by the decreased cilia length in ALI organotypic cultures in which Notch was inactivated in Rbpjk f/f adult progenitors transduced using a lentivirus-expressing Cre approach (Fig. S8).

Moreover Notch-Prom1 interactions have been reported in various cellular contexts: Notch-signaling has been shown to positively regulate Prom1-expression in human breast cancer cells (42), and in different types of gastric cancer cells, specifically through an RBPJκ dependent-pathway (22). Given that Notch-signaling is known to prevent the differentiation of airway progenitors into multiciliated cells, we reason that Notch-activation in these cells occurs after the commitment to the multiciliated cell lineage, when even low levels of Notch-signaling may be sufficient to modulate the expression of cilia-related genes and consequently fine tune the ciliary phenotype.

Notably, a recent comprehensive single cell RNAseq-based survey of the cellular components of the human lung identified two regionally distinct populations of multiciliated cells that differ in their location and molecular signature: “proximal ciliated”, and “ciliated”. Further analysis of the published database (Synapse [https://www.synapse.org/#!Synapse:syn21041850]; cellxgene at https://hlca.ds.czbiohub.org) revealed that these populations differentially express Notch receptors (NOTCH1, NOTCH2) and the Notch target HES1, all of which are particularly enriched in the multiciliated “non-proximal” cell population (17). Therefore, as also suggested by our study, multiciliated cells differ in their ability to express and activate Notch. We hypothesize that this is likely to reflect in regional differences in Prom1 expression and ultimately in differences in the multiciliated cell phenotype along the respiratory tract.

We propose that Prom1 and Notch-signaling are integral components of a mechanism responsible for maintaining the proper structure and function of motile cilia in the multiciliated cells throughout the airway epithelium. Our observations of the graded Prom1 expression in vivo suggest that Prom1 is kept at different levels regionally, likely in response to local signals, to prevent inappropriate cilia growth or ciliary beating. This mechanism contributes to generate distinct features, such as ciliary length and dynamics in multiciliated cells from different airway generations, thus resulting in diversity of this cell type. In this regard our model predicts that increases in cilia length by transient activation of Notch signaling would trigger a mechanism to restrict cilia growth in which Prom1 has a critical role (Fig. S9). In our Prom1KO cultures ciliary growth, presumably by Notch and other factors was unrepressed, resulting in longer cilia. A similar mechanism could be invoked for regulation of ciliary dynamics

We argue that this mechanism is also likely to be relevant during repair of damaged shortened cilia from exposure to cigarette smoke and biological agents, as well as in chronic pulmonary conditions, such as COPD (43). Lastly, we also consider that the mechanisms described here may be relevant in the context of multiciliated cells in other mammalian organs, including that of the reproductive tract.

## Materials and Methods

### Immunofluorescence (IF)

Whole E14.5 embryos and E18.5 lungs and tracheas were fixed overnight at 4ºC in 4% paraformaldehyde, processed for frozen OCT embedding and sectioned (5um) in a cryotome, as previously reported (44). Transwell membranes of ALI airway organotypic cultures were fixed with 4% paraformaldehyde for 10 minutes at room temperature and processed as described (45). For IF, tissue sections were incubated overnight at 4 ºC with primary antibody, while for Transwell membranes antibody incubation was performed at room temperature for 2 hours. For all samples, IF signals were detected using Alexa Fluor 488, Alexa Fluor 568, or Alexa 647-labeled secondary antibodies (1:250) (Thermo Fisher, Waltham, MA) for 2 hours at room temperature. 4′,6-diamidino-2-phenylindole (DAPI) was used to stain intracellular DNA (NucBlue, Thermo Fisher, Waltham, MA). Alexa Fluor 647 Phalloidin (Thermo Fisher, Waltham, MA) was used to visualize filamentous actin (F-actin). Following PBS washes, tissue sections and membranes were mounted with anti-fade mounting media (ProLong Gold, Thermo Fisher, Waltham, MA) and examined. The antibodies used were: anti-CD133 (Prominin-1) (Thermo Fisher #14-1331-82, 1:100), anti-Foxj1 (Thermo Fisher #14-9965-80, 1:100), anti-Acetyl-α-Tubulin (Cell Signaling #5335, 1:300), anti-beta IV Tubulin (Abcam #ab11315, 1:100), anti-Cetn3 (Abnova #H00001070-M01, 1:50), anti-p63-α (Cell Signaling #13109, 1:100), anti-RBPSUH (Cell Signaling #5313, 1:100), anti-UGRP1 (R&D Systems, #MAB3465, 1:100), and anti-Cc10 (Santa Cruz Biotechnology #sc-9772, 1:100). Confocal laser scanning micrographs were acquired using a Zeiss LSM-710 laser scanning microscope (Zeiss, Oberkochen, Germany) equipped with a motorized stage, 20x dry or 63x oil-immersion objectives, and an argon laser. For Z-stack analysis, the Z-stepper was configured to take 130 nm steps. The remaining fluorescent micrographs were taken with a Leica DMi8 widefield fluorescence microscope (Leica Microsystems, Wetzlar, Germany), and a 40x dry objective.

### Cytometry and Fluorescence Intensity Analyses

For qualitative assessment of Prom1 fluorescence intensity *in vivo*, we performed Prom1 immunostaining of E18.5 tracheas and lungs from control and Rbpjκ^cnull^ mutant mice. The intensity of fluorescence detection was evaluated in single fields representative of different airway generations (trachea, primary bronchus, secondary bronchus and bronchiole) by setting a threshold that limited the ROI to the areas where Prom1 signal was detected, and then calculating the mean gray value and its standard deviation with the Fiji software (46). The resulting measurements were normalized as a percentage to the most proximal or distal value, whichever was highest, and plotted on a bar chart with Prism 8.4.3 software (GraphPad). For qualitative assessment of Prom1 fluorescence intensity *in vitro*, we performed Prom1 immunostaining and F-actin labeling with Phalloidin in ALI day 0 and ALI day 8 mouse airway epithelial cell cultures. The samples were obtained from mouse tracheal epithelial cells that had been incubated with the gamma-secretase inhibitor DAPT, or DMSO for vehicle controls, since 3 days prior to confluence being reached (ALI day -3). The intensity of fluorescence detection was evaluated in single representative fields by delimiting the cell borders evidenced by Phalloidin with the Polygon tool, and calculating the mean gray value of Prom1 detection in each Prom1-labelled individual cell with the Fiji software (46). The resulting measurements were plotted on vertical violin plots. The frequency distributions of intensity values (with a bin width of 5) were plotted on bar charts, and curve-fitted with non-linear regression (Gaussian or sum of two Gaussians).

All the analyses and plots were done with Prism 8.4.3 software (GraphPad). For quantitative assessment of the number of Foxj1-positive cells *in vitro*, we performed Foxj1 immunostaining of ALI day 8 mouse airway epithelial cell cultures. The samples were obtained from mouse tracheal epithelial cells that had been transduced with a lentivirus containing either an sgRNA sequence targeting Foxj1 (Foxj1 KO), or a scrambled sgRNA sequence (control). The number of Foxj1-positive cells was measured in 10 random fields for each condition by using the Multi-point tool of the Fiji software (46). The individual values were plotted in a beeswarm plot with the sample-mean and respective standard deviation superimposed. The sample-means were compared with a two-tailed unpaired t-test. All the analyses and plots were done with Prism 8.4.3 software (GraphPad). For quantitative assessment of the number of p63-positive cells *in vitro*, we performed p63 immunostaining of ALI day 0 mouse airway epithelial cell cultures. The samples were obtained from mouse tracheal epithelial cells that had been transduced with a lentivirus containing either an sgRNA sequence targeting Prom1 (Prom1 KO), or a scrambled sgRNA sequence (control). The number of p63-positive cells was measured in 20 random fields from 3 independent ALI culture wells for each condition, by using the Multi-point tool of the Fiji software (46). The individual values and the well-means were plotted as a beeswarm plot with the sample-mean and respective standard deviation superimposed. The sample-means were compared with a two-tailed unpaired t-test. All the analyses and plots were done with Prism 8.4.3 software (GraphPad). For quantitative assessment of the number of Cetn3-positive cells *in vitro*, we performed Cetn3 immunostaining of ALI day 4 mouse airway epithelial cell cultures. The samples were obtained from mouse tracheal epithelial cells that had been transduced with a lentivirus containing either an sgRNA sequence targeting Prom1 (Prom1 KO), or a scrambled sgRNA sequence (control). The number of Cetn3-positive cells was measured in 20 random fields from 3 independent ALI culture wells for each condition, by using the Analyze Particles tool of the Fiji software (46). The individual values and the well-means were plotted as a beeswarm plot with the sample-mean and respective standard deviation superimposed. The sample means were compared with a two-tailed unpaired t-test. All the analyses and plots were done with Prism 8.4.3 software (GraphPad).

### In situ hybridization

In situ hybridization was performed in frozen sections as we previously described in Tsao *et al* (7). The hybridization probe was synthesized from a Mammalian Gene Collection (MGC) fully sequenced mouse Prom1 cDNA (4502359, Dharmacon, Lafayette, CO). After confirmation by Sanger sequencing, the plasmid was linearized, and then amplified with Q5 High-fidelity DNA polymerase (M0491S, New England BioLabs, Ipswich, MA) according to the manufacturer’s instructions. The forward primer used was AATTAACCCTCACTAAAGGGCTTTGAGTGAATGACCACCTTG and the reverse primer was TAATACGACTCACTATAGGGGCCTTGGAATCAACTGAGATGTC. The PCR product was transcribed with QIAquick PCR Purification Kit (#28104, Qiagen, Hilden, Germany), transcribed with MAXIscript Kit (#AM1320, Thermo Fisher, Waltham, MA), and purified with RNeasy MinElute Cleanup Kit (#74204, Qiagen, Hilden, Germany), according to manufacturer’s instructions.

### Mouse models

Rbpjk^cnull^ and ShhCre;NICD mice were generated as described by Tsao *et al* (7) and Guha *et al* (29). The detection of vaginal plugs in the morning was considered as day 0.5, and embryos were harvested at either E14.5 or E18.5.

### Air-liquid-interface (ALI) organotypic airway epithelial cultures

Mouse tracheal epithelial cells were isolated from the tracheas of adult (8-12 weeks old) C56BL/6J mice and released from tissue by pronase digestion (1.5 mg/mL) for 24 hours at 4ºC. Airway epithelial cells were isolated from fibroblasts by differential adhesion, as previously described (27). Freshly isolated mTEC were seeded at a density of 1×10^5^/cm^2^ on supported semi-permeable polyester membranes (Transwell, Corning-Costar, Corning, NY) that had previously been coated with collagen. The medium used to initiate the cell culture was mTEC/Plus supplemented with 10% FBS and retinoic acid. Once the cells were confluent (Day 0), and to induce their differentiation, the medium from the upper chamber was aspirated, thereby inducing an air-liquid-interface (ALI), and the medium from the bottom chamber was replaced with mTEC/Serum-Free (mTEC/Basic) supplemented with retinoic acid, as previously described (27). Cell cultures were maintained by replacing the respective media every 48h or less, for up to 27 days after inducing the air-liquid interface. For genetic disruption of Notch signaling in mTEC ALI cultures, adult basal cells from *Rbpjk*^*flox/flox*^ mice were isolated and transduced with a lentivirus-expressing Cre as we previously reported (30). For Notch pharmacological inhibition, mTEC were cultured from ALI day -3 to ALI day 8 with an added gamma-secretase inhibitor (DAPT, 50 μM, Sigma) or vehicle control (DMSO, Sigma, D8418), as previously described.

### Plasmid construction, lentivirus production and lentiviral transduction

3 sgRNAs targeting the Prom1 gene and 3 sgRNAs targeting the Foxj1 gene were designed and selected according to their predicted sgRNA efficient scores, as calculated by the sgRNA designer tool provided by Broad Institute (available at: http://www.broadinstitute.org/rnai/public/analysis-tools/sgrna-design-v1). A scrambled-sequence sgRNA was designed to serve as a negative control. The sgRNA oligos were cloned into lentiCRISPRv2 plasmid (Addgene) as previously described in the Target Guide Sequence Cloning Protocol from the Zhang Lab (available at: https://www.addgene.org/crispr/reference/#protocols). Each plasmid containing an inserted sgRNA sequence was verified using Sanger sequencing. The VSV-G pseudotyped lentivirus was packaged by 5 plasmid co-transfection of HEK293T cells using TansIT Transfection Reagent (Mirus Bio) in 15 cm culture dishes (The protocol describing production of lentiviral particles can be found at http://www.www.bu.edu/dbin/stemcells/protocols). HEK293T cells were cultured in high glucose DMEM medium containing 10% FBS. The supernatant was collected for a total of 5 times, starting at 48 hours after transfection and every 12 hours afterwards, and readily filtered with a 0.45 μm pore size vacuum filter. The filtrate was concentrated by spinning for 90 minutes at 48,000 g, resuspended in high glucose DMEM medium, and stored at -80°C. For lentiviral transduction, the protocol outlined by Horani *et al* was used (32). Following the isolation by differential adhesion, mouse airway epithelial cells were resuspended in mTEC/Plus supplemented with retinoic acid, Polybrene, and 10 mM Y27632. At this point, 10 μL of concentrated lentivirus filtrate were added to a 1.5 mL microfuge tube containing 150 μL of resuspended cells. After 10 minutes of incubation at room temperature, the cells were delivered to the apical surface of the supported membranes. After 24 hours of incubation at 37ºC, the culture medium was aspirated from the bottom chamber only, and replaced with fresh mTEC/Plus supplemented with retinoic acid and Y27632. In order to maximize adhesion, the medium on the upper chamber was not aspirated for the first 48 hours after transduction, at which time it was replaced with fresh mTEC/Plus supplemented with retinoic acid and Y27632. Starting 72 hours after the initial exposure to the lentivirus, the culture medium was supplemented with 2.0 μg/mL puromycin for two consecutive 48-hour periods, resulting in a total of 4 days of puromycin treatment. The culture medium (mTEC/Plus) was supplemented with Y27632 for the entire duration of the submerged conditions, which was only discontinued when the medium from the upper chamber was aspirated and the air-liquid interface induced (ALI day 0). The sgRNA with the highest expression-disrupting efficiency for each gene, as assessed by cytometric analysis of immunofluorescence for Prom1 at ALI day 0, or Foxj1 at ALI days 4 and 8, was used in subsequent experiments. The sgRNA sequence used for the targeted disruption of Prom1 was TGAGCAGACAAATCACCAGG. The sgRNA sequence used for the targeted disruption of Foxj1 was TGGGGGTCTGTGCCTCCTGG. Generation of Lentivirus-expressing Cre and expression in ALI cultures were previously reported (30).

### Real-time PCR/transcriptome analyses

For quantitative real-time PCR analysis of the organotypic airway cultures Transwel membranes containing mTEC-derived cells were cut from the supporting structure and submerged in RLT lysis buffer. RNA was extracted with an RNeasy Mini kit (#74106, Qiagen, Hilden, Germany), according to the manufacturer’s instructions. First-strand cDNA was synthesized with the SuperScript IV First-Strand Synthesis System (#18091050, Thermo Fisher, Waltham, MA), according to the manufacturer’s instructions. Quantitative RT-PCR was performed as previously described by using Taq-Man Advanced Master Mix (#4444556, Thermo Fisher, Waltham, MA) and a StepOnePlus™ Real-Time PCR System (Applied Biosystems, Foster City, CA). The following primers (Thermo Fisher) were used: Prom1 (Mm01211402_m1), Foxj1 (Mm01267279_m1), Scgb3a2 (Mm00504412_m1) and Cetn3 (Mm00514305_m1). ACTB was used as an internal control. The changes in expression levels were calculated using the ΔΔCt method. The results were either plotted as Relative Expression Levels (Fold Change ± CI95) in a bar chart, or as ΔCt (CtTarget-CtACTB) in a beeswarm plot with the mean ± SD superimposed, both with Prism 8.4.3 software (GraphPad). Regarding the latter, a lower ΔCt corresponds to decreased expression of the target gene. Transcriptome analysis was previously performed in E18.5 WT, Rbpjk^cnull^, ShhCre;NICD*1*(7)(29)(44) and deposited (GEO, Series ID (GSE52926).

### High Speed Video Microscopy

Mouse tracheal epithelial cells cultured *in vitro* on supported membranes were imaged live with a 40x objective on a live cell-configured inverted microscope (Axio Observer Z1 platform equipped with ApoTome.2, Zeiss, Oberkochen, Germany) attached to a high-speed video camera in order to capture an *en face* view of distal tips of beating cilia. Videos were captured in 10 random locations in each membrane.ALI cultures were also scraped so that clusters of ciliated cells could be placed on a microscope slide and the cilia viewed from the side. To achieve this, 20 μL of culture medium were added to the surface of the ALI cultures, a small area of cells scraped with a 200 μL micropipette-tip, and the cells suspended in medium and transferred to a microscope slide, as previously described (37). All cells were imaged inside a live cell-imaging chamber heated to a stable temperature of 37°C, at a frame rate of 360 frames per second, and for a duration of at least 720 frames using CoreView (v2.2.0.9) software (IO Industries, London, ON, Canada). Mucociliary transport was assessed by tracking the displacement of microspheres added to the apical surface of live ALI cultures. The cultures were imaged under conditions similar to the above, but with a 10x objective, a fluorescent light source, and at a frame rate of 25 frames per second, as previously described (37). Videos were captured in 5 random locations in each membrane.

### Ciliary beating frequency, Cilia length and Mucociliary transport analyses

Ciliary Beating Frequency (CBF) was calculated from *en face* video recordings of mTEC cultured on supported membranes and of scraped multiciliated cells placed on microscope slides. CBF was calculated for each video by using the fast Fourier transform (FFT) in Matlab (R2019a, Mathworks, Natick, MA), as previously described (37). In order to quantify the CBF across each video frame, the frequency of the highest magnitude peak in the FFT was identified for each pixel. Only pixels in which the FFT indicated a magnitude greater than 1, in units of pixel intensity, and frequency greater than or equal to 2 Hz were included in the CBF heatmap and used to calculate the video average, as originally described by Sisson et al. (47).The averages from each video and the well-means were plotted as a beeswarm plot with the sample-mean and respective standard deviation superimposed. The sample-means were compared by using a two-tailed unpaired t-test, or by using ordinary one-way ANOVA followed by Dunnett’s multiple comparison tests (alpha 0.05). All the analyses and plots were done with Prism 8.4.3 software (GraphPad). Cilia length (CL) was assessed in adult mTEC cultures in multiciliated cells scraped from the membranes and placed on microscope slides. Damaged cilia, or those in close proximity to other cells and/or debris were not excluded. Five individual cilia per multiciliated cell were measured and the average cilia length per cell was determined. Length values (in pixels) were normalized so that the average cilia length of the control sample equaled 1. The well-means were plotted as a beeswarm plot superimposed on a bar chart representing the sample-means and respective standard deviations. The means were compared using a two-tailed unpaired t-test (Prism 8.4.3 software, GraphPad). In the experiments involving lenti-Cre transduction of WT or Rbpjκ^f/f^ ALI cultures CL was assessed in ten individual cilia from a representative confocal laser scanning micrograph of each condition using Zeiss LSM imager browser. Sample-means were plotted in a bar chart with their respective standard deviations and differences were compared using a two-tailed unpaired t-test. In a subset of multiciliated cells from control ALI cultures CL and CBF were analyzed simultaneously. The correlation between these two variables was analyzed by computing Pearson correlation coefficients (two-tailed, alpha of 0.05) using a simple linear regression. The CL and CBF pairs were plotted in a scatter plot with the linear regression line (solid) and CI 95 (dashed) superimposed. Mucociliary Transport (MCT) was assessed by automatically tracking the displacement of fluorescent microspheres (1 μm diameter, Fluoro Max Dyed Green, G0100, ThermoFisher, Waltham, MA) added to the surface of live ALI cultures by using the Fiji plugin TrackMate (38). The solution of microspheres was diluted 1:1,000 in PBS, approximately 15 μL of the diluted microsphere solution was added to the surface of the ALI, and the culture plates returned to the incubator for approximately 10-40 minutes prior to imaging, as previously described (37). The final MCT value was calculated by weight-averaging the track displacement velocity by the track displacement distance, thereby accounting for microspheres that may have gone in- and out-of-focus as well as for microspheres that may have only briefly crossed the microscope field. The averages from each video and the well-means were plotted as a beeswarm plot with the sample-mean and respective standard deviation superimposed. The sample-means were compared with a two-tailed unpaired t-test. All the analyses and plots were done with Prism 8.4.3 software (GraphPad).

### Immunoblotting

For immunoblotting analysis of Prom1, mouse airway epithelial cells grown on permeable supported membranes were washed at ALI day 0 with cold PBS and scraped off. The samples were obtained from the proliferation of mouse tracheal epithelial cells that had been transduced with a lentivirus containing either an sgRNA sequence targeting Prom1 (Prom1 KO) or a scrambled sgRNA sequence (control). Proteins were analyzed by SDS-polyacrylamide gel electrophoresis (SDS-PAGE, 12%), and transferred to Immobilon-P PVDF Membrane (IPVH00010, Millipore, Belfore, MA), according to standard procedures. Immunoblotting was performed as previously described by Corbeil et al (48), using anti-CD133 (Prominin-1) (Thermo Fisher #14-1331-82) as a primary antibody.

## Acknowledgments

We thank Steve Brody for the help with the HSVM script. Kyle Travaglini and Mark Krasnow for their help in the analysis of the multiciliated cell transcriptome from their single cell RNAseq database. We also thank members of the Cardoso and Lu labs at CCHD for the helpful discussions. All diagrams/illustrations in the manuscript were created with BioRender.com. This work was supported by NIH-NHLBI R35-HL135834-01 to W.V.C and Fundação para Ciência e a Tecnologia MD PhD Scholarship PD/BD/113766/2015 to C. F. H. S.

## Supplementary Information

**Figure S1.**
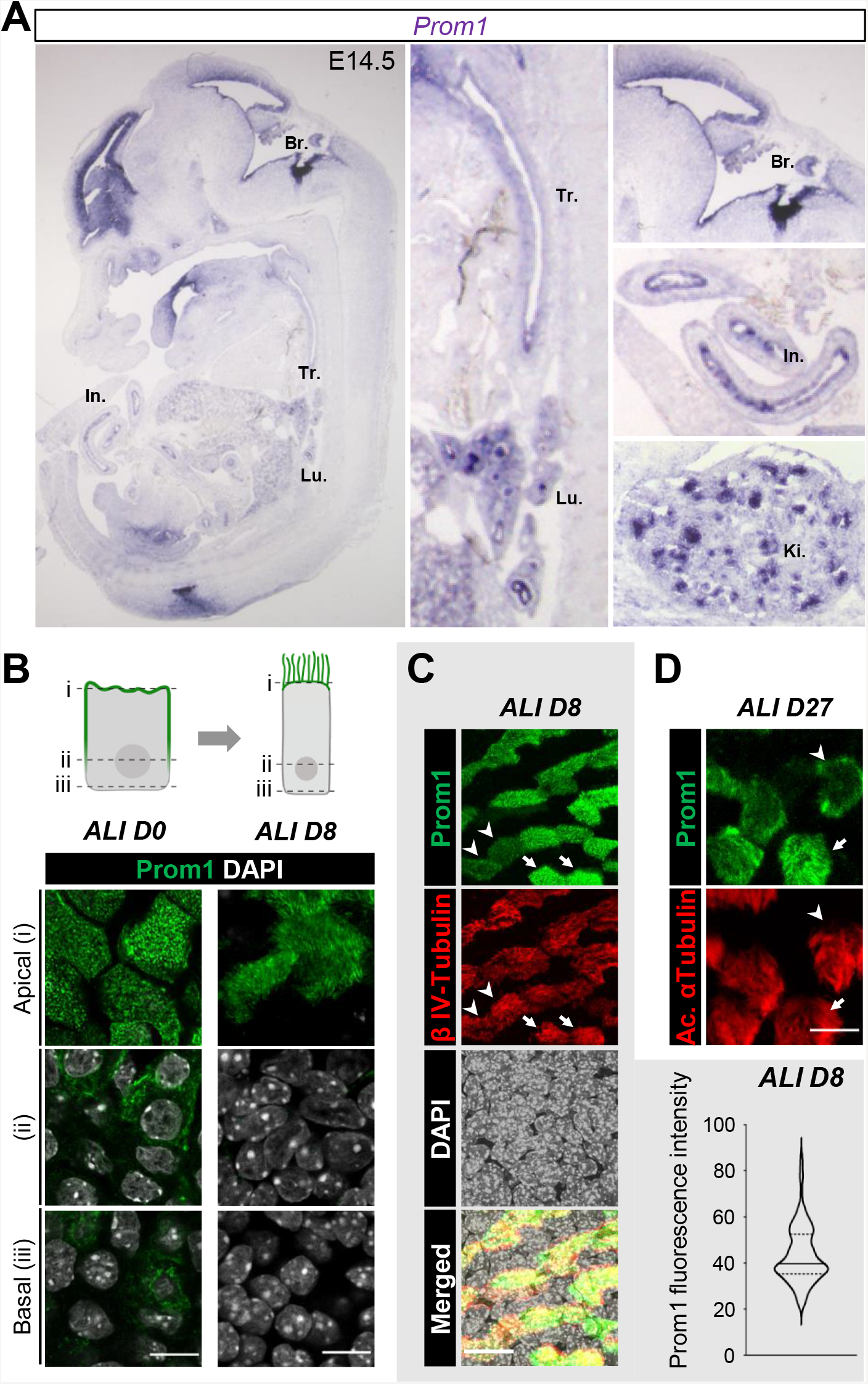
Prom1 is broadly expressed in epithelial progenitors of developing organs and in differentiating adult airway progenitors. **(A)** *In situ* hybridization of Prom1 in E14.5 wild type embryos. Prom1 expression in lung (Lu.), trachea (Tr.), brain ependyma-choroid plexus (Br.), intestine (In.), and kidney (Ki.). **(B)** Immunofluorescence (IF) of Prom1 in air-liquid interface (ALI) cultures at days 0 and 8. Each panel represents a series of single optical sections along the apical-basal axis of cells as depicted in the diagram. Prom1 labelling is largely restricted to the apical domain of both the undifferentiated monolayer at day 0 and the pseudostratified epithelium at day 8, with light labelling also present in the basolateral compartment at day 0. **(C)** Double IF of Prom1 and β IV-tubulin in ALI cultures at day 8. Prom1 labelling is found selectively in multiciliated cells and with wide range of signal intensities (arrowhead, weak; arrow, strong) which reflect the heterogeneity among these cells. The graph represents a vertical violin plot of the intensity of Prom1-labelling in individual Prom1+ cells from a representative field of an ALI culture at day 8. The solid line represents the median and the dashed lines represent the quartiles. The values represent pixel intensity from 0 to maximal (arbitrary units). **(D)** Double IF of Prom1 and acetylated α-tubulin in ALI cultures at day 27. Prom1 labelling is still found selectively in multiciliated cells and with a wide range of signal intensities (arrowhead, weak; arrow, strong) reflecting heterogeneity of expression. DAPI: gray in all panels. Scale bars in B, C and D = 10 μm.

**Figure S2.**
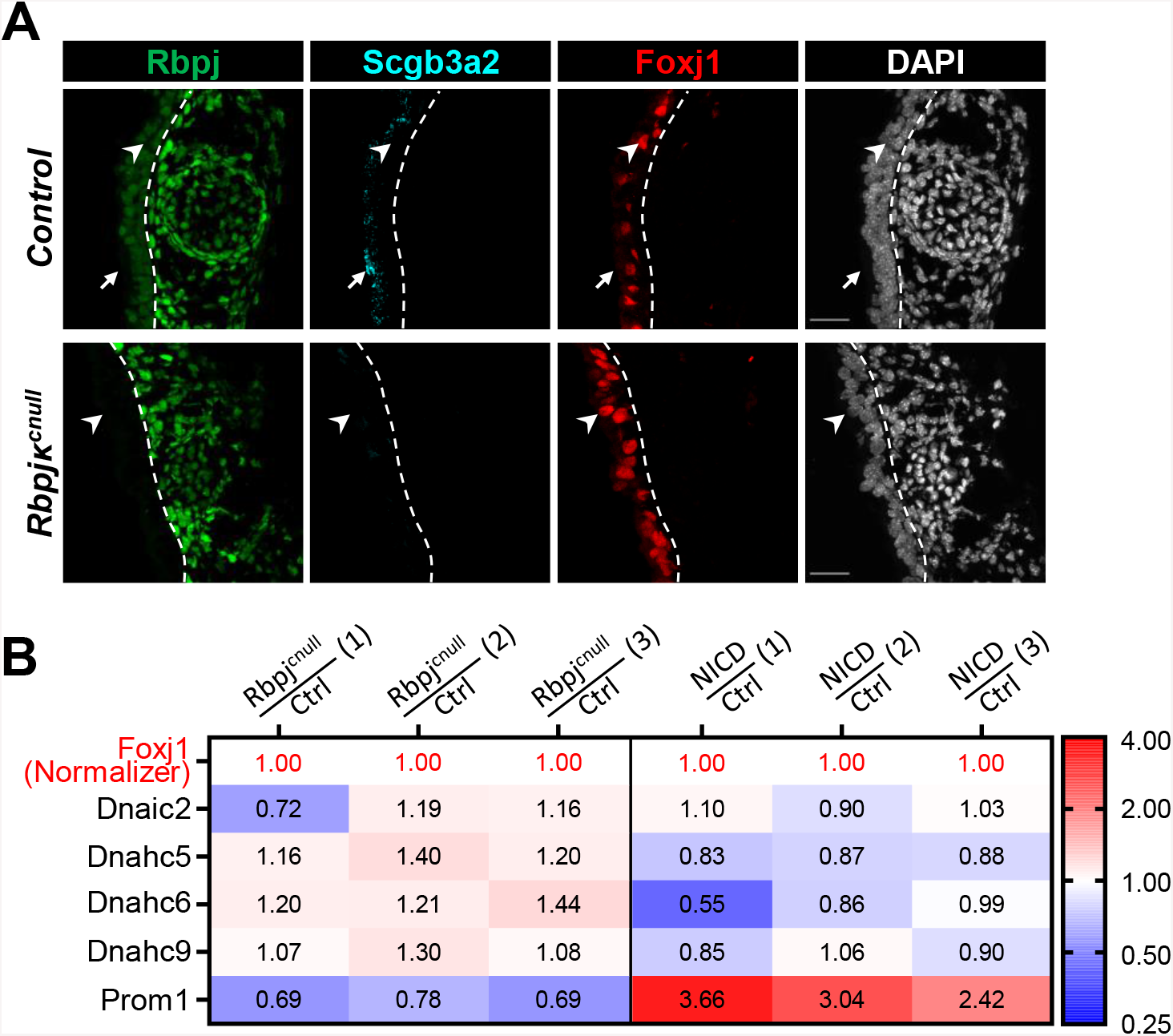
Effects of Notch gain or loss of function in the developing airway epithelium. **(A)** Triple immunofluorescence of Rbpjκ, Scgb3a2 and Foxj1 in E18.5 wild type and Notch-deficient (Rbpjκ^cnull^) tracheas. The dashed line demarcates the basement membrane. The arrows point to a secretory cell (labelled by Scgb3a2) and the arrowheads point to a multiciliated cell (labelled by Foxj1). In the Notch-deficient airways there is no expression of Rbpj in the epithelium (left of the dashed line), an overpopulation of multiciliated cells, and absence of secretory cells. Scale bars = = 25 μm. **(B)** Expression of cilia-associated genes (including Prom1) in the lung epithelium of E18.5 control (WT), Rbpjκ^cnul^ and ShhCre;NICD mice from whole lung transcriptome analysis (n=3 per group). Values are normalized by Foxj1 and expressed as relative fold change from Foxj1 values. Prom1 is consistently downregulated in all Rbpjκ^cnull^ samples and upregulated in ShhCre;NICD.

**Figure S3.**
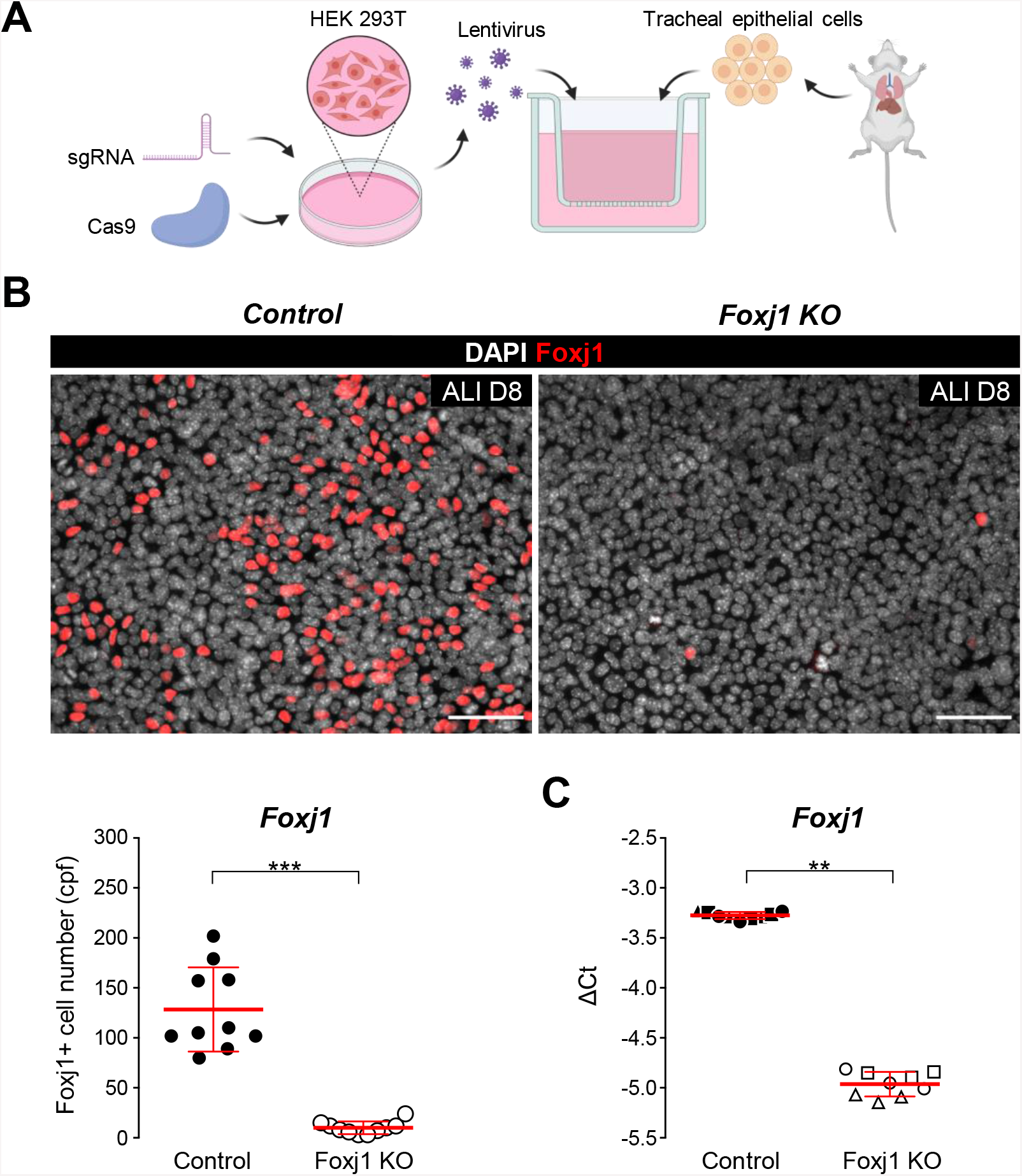
CRISPR/Cas9-mediated disruption of Foxj1-expression in organotypic ALI cultures. **(A)** Diagram of experimental strategy: sgRNA targeting of Foxj1, lentivirus production and transduction of mTECs (see methods). **(B)** Top panels: representative micrographs of immunofluorescence of Foxj1 in ALI day 8 cultures incubated with CRISPR/Cas9 lentiviral vectors carrying a scrambled-sequence (Control, left) or a Foxj1-targeting sequence (Foxj1 KO, right). Bottom panel: cytometric analysis of the number of Foxj1-labelled cells in ALI day 8 control and Foxj1 KO cultures; the beeswarm plot represents the number of Foxj1+ cells per field (cpf) in 10 random fields (●, N=10), with the well-mean ± SD superimposed in red. In Foxj1 KO cultures there is a dramatic decrease in the number of Foxj1+ cells. Student’s t-test: ***, *p* < 0.001 **(C)** qPCR analysis of Foxj1 in Control and Foxj1 KO day 8 ALI cultures showing significant decrease Foxj1 expression in Foxj1 KO cultures compared to control. The graph represents individual ΔCt values (N_technical replicates_ = 3, N_wells_ = 3), with the sample mean ± SD superimposed in red. ACTB was used as the reference gene. Student’s t-test: **, *p* < 0.01. DAPI is gray in all panels. Scale bars in B = 50 μm

**Figure S4.**
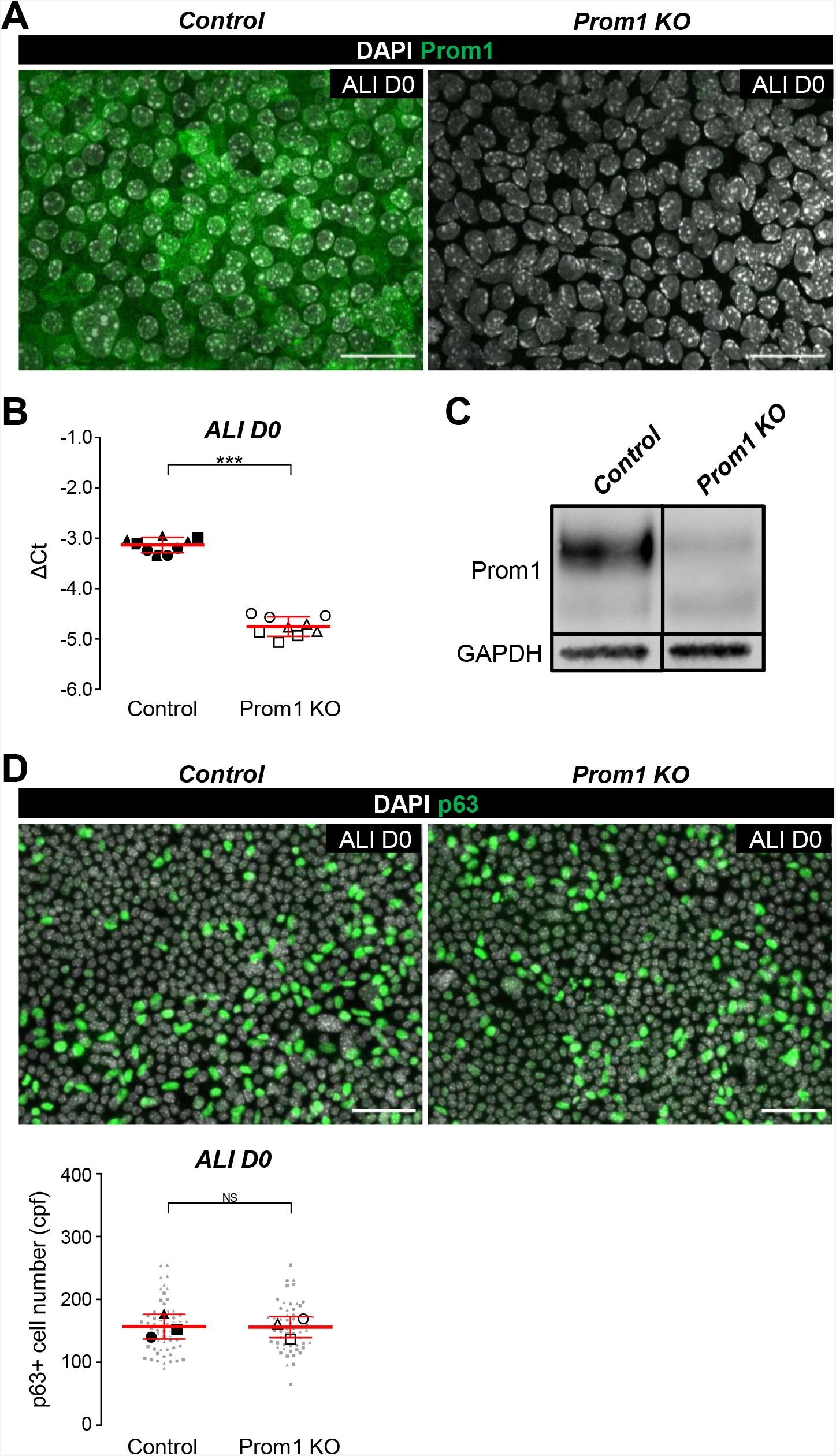
Prom1 inactivation does not prevent expansion of airway epithelial progenitors. **(A)** Representative micrographs of Prom1 immunofluorescence (IF) in airway progenitors (mTEC ALI day 0 cultures) incubated with CRISPR/Cas9 lentiviral vectors carrying a scrambled-sequence (Control, left) or a Prom1-targeting sequence (Prom1 KO, right). In Prom1 KO cultures there is no detectable labelling of Prom1. **(B)** QPCR analysis of Prom1 in Control and Prom1 KO day 0 ALI cultures: significant decrease in Prom1 expression in Prom1 KO cultures compared to control. The graph represents individual ΔCt values (N_technical replicates_ = 3, N_wells_ = 3), with the sample mean ± SD superimposed in red. ACTB was used as the reference gene. Student’s t-test: ***, *p* < 0.001. **(C)** Representative Western Blot analysis of Prom1 in Control and Prom1 KO day 0 ALI cultures (N = 3). There is a significant decrease in Prom1 expression in Prom1 KO cultures compared to control. GAPDH was used as a loading control. **(D)** Top panels: representative p63 IF in ALI day 0 cultures incubated with CRISPR/Cas9 lentiviral vectors carrying a scrambled-sequence (Control, left) or a Prom1-targeting sequence (Prom1 KO, right). Bottom panel: cytometric analysis of the number of p63-labelled cells in ALI day 0 control and Prom1 KO cultures; the beeswarm plot represents the number of Prom1+ cells per field (cpf) in 20 random fields per well (•, N=20), in a total of 3 wells per condition (●, N = 3), with the sample-mean ± SD superimposed in red. No significant (NS) difference between the number of p63+ cells in Prom1 KO cultures compared to control (Student’s t-test).

**Figure S5.**
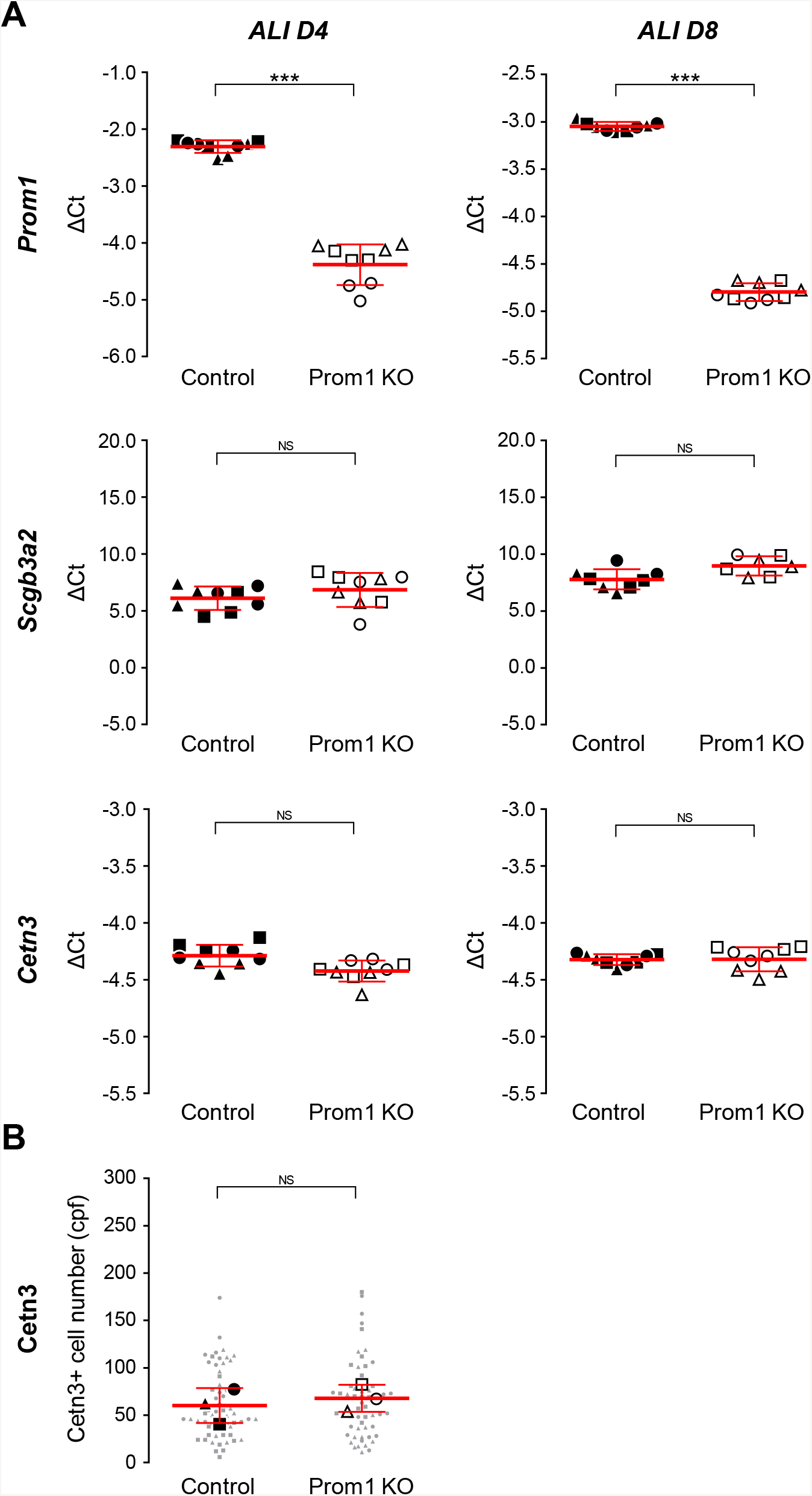
Prom1 inactivation does not alter the balance of ciliated *vs*. secretory phenotype in the airway epithelium. **(A)** QPCR analysis of Prom1, Scgb3a2 and Centrin3 in Control and Prom1 KO ALI cultures at days 4 and 8. There is a significant decrease in the expression of Prom1 but not of Scgb3a2 or Centrin 3 in Prom1 KO cultures compared to control at these stages. Graphs represent individual ΔCt values (N_technical replicates_ = 3, N_wells_ = 3), with the sample mean ± SD superimposed in red. ACTB was used as the reference gene. Student’s t-test: NS, not significant; ***, *p* < 0.001. **(B)** Cytometric analysis of the number of Centrin3-labelled cells in ALI day 4 control and Prom1 KO cultures; the beeswarm plot represents the number of Centrin3+ cells per field (cpf) in 20 random fields per well (•, N=20), in a total of 3 wells per condition (●, N = 3), with the sample-mean ± SD superimposed in red. No significant (NS) difference between the number of Centrin3+ cells in Prom1 KO cultures when compared to control (Student’s t-test).

**Figure S6.**
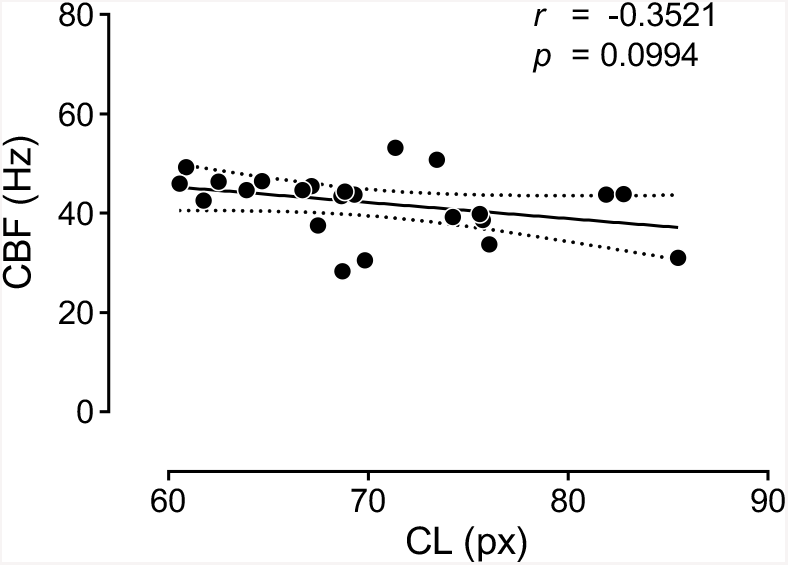
Cilia Length (CL) and Ciliary Beating Frequency (CBF) are not correlated in multiciliated cells from ALI cultures. Individual multiciliated cells scraped from control mTEC ALI cultures at day 27 were analyzed for both CL and CBF. The correlation between these variables was determined by Pearson correlation coefficients (two-tailed, alpha of 0.05) and by using a simple linear regression. The graph represents individual multiciliated cells plotted according to their average CBF (Hz) and CL (px, pixels), with a simple linear regression (solid line, Y = -0.3236*X + 64.82) ± CI 95 (dashed line) superimposed. In multiciliated cells from control ALI, CBF and CL are not correlated Pearson correlation: r = -0.352; *p* = 0.0994.

**Figure S7.**
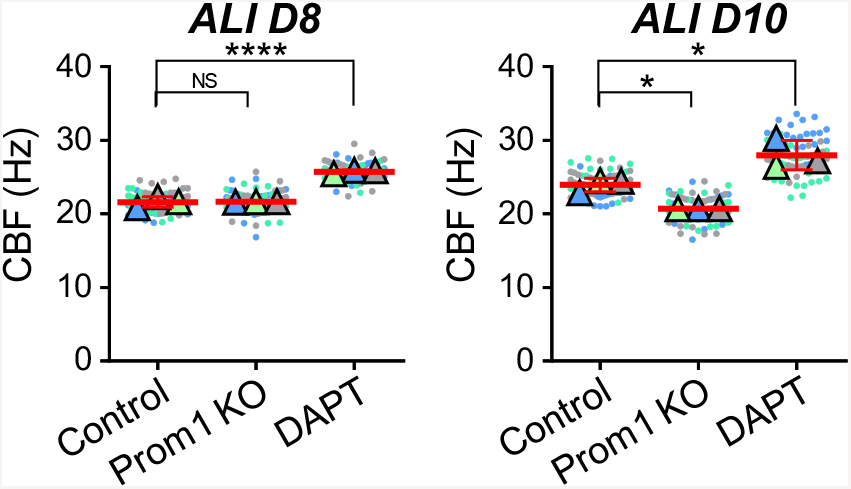
DAPT effects in CBF precedes that of Prom1 KO in differentiating ALI cultures. Analysis of Ciliary Beating Frequency (CBF) in ALI cultures incubated with CRISPR/Cas9 Control or Prom1 KO lentivirus, and with the Notch-inhibitor (DAPT) or its vehicle control (DMSO), at ALI days 8 and 10. The graphs are beeswarm plots of the average CBF, in Hz, of each video (●, N = 25) and well (Δ, N = 3), with the sample-means ± SD superimposed in red. Notch-deficient cultures have increased CBF when compared to controls at both stages. By contrast CBF is not significantly affected in Prom1 KO at ALI day8 compared to control and is found decreased only later at ALI day 10. Dunnett’s multiple comparison tests: ns, not significant; *, adjusted *p* value < 0.05; ****, adjusted *p* value < 0.0001.

**Figure S8.**
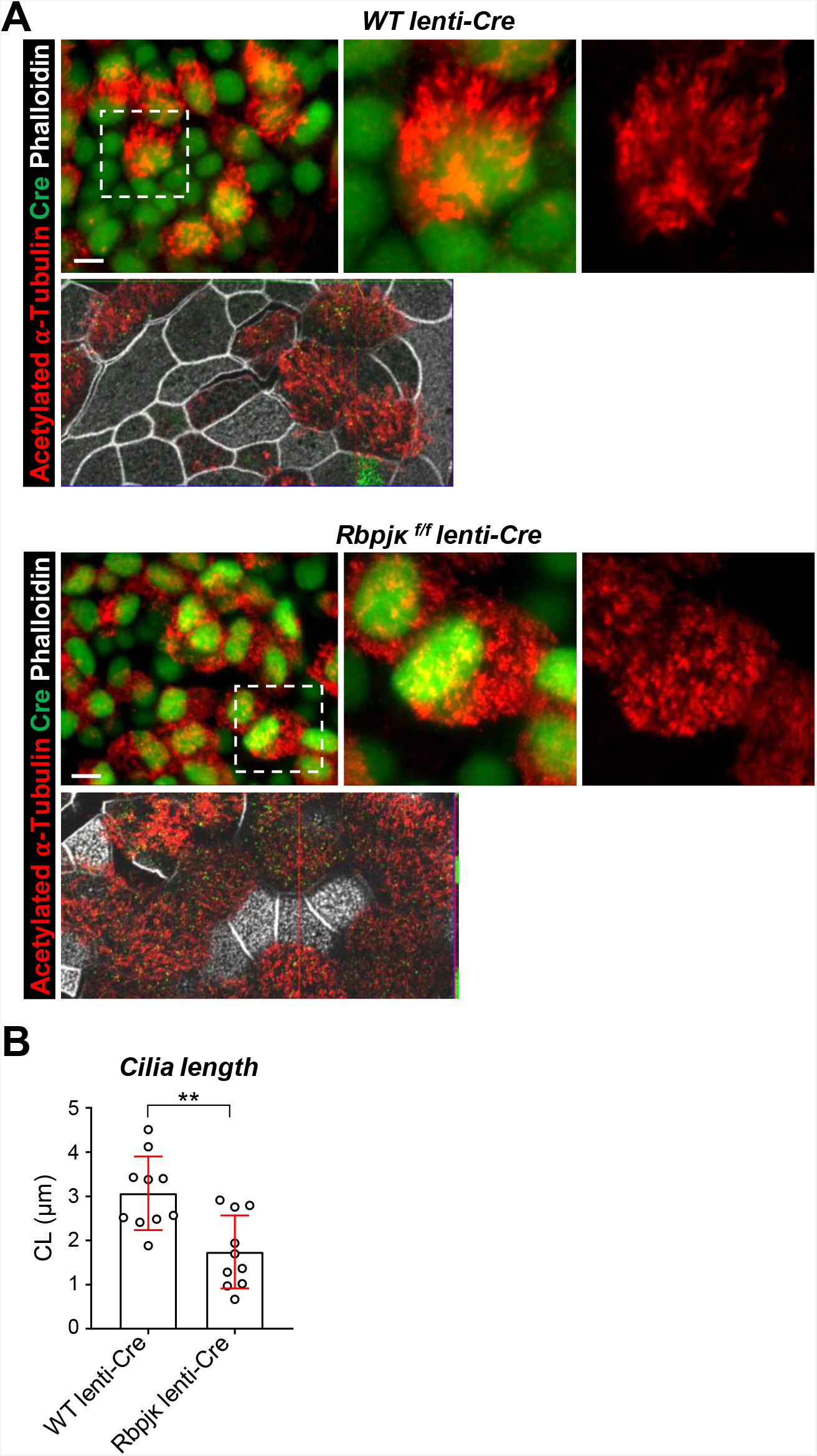
Decreased cilia length (CL) in multiciliated cells from Rbpjκ null ALI cultures. **(A)** Double immunofluorescence of acetylated α-Tubulin and Cre in mTEC ALI cultures from airway progenitors of WT or Rbpjκ^f/f^ adult mice, incubated with Ef1α-Cre lentivirus at the time of plating. Disruption of Notch in Rbpjκ^f/f^ lenti-Cre leading to aberrant expansion of the multiciliated cells (as and shortened cilia (boxed areas enlarged in right panels, phalloidin in gray, Bars = 8 um). **(B)** Quantitative analysis of CL in WT lenti-Cr and Rbpjκ^f/f^ lenti-Cre ALI cultures. Graph CL measurements of 10 single cilia (N_cilia_ = 10) from random fields in each condition (Bar is mean ±SE). CL significantly reduced in Rbpjκ^f/f^ lenti-Cre cultures compared to WT-lenti-Cre. Student’s t-test: **, *p* < 0.01.

**Figure S9.**
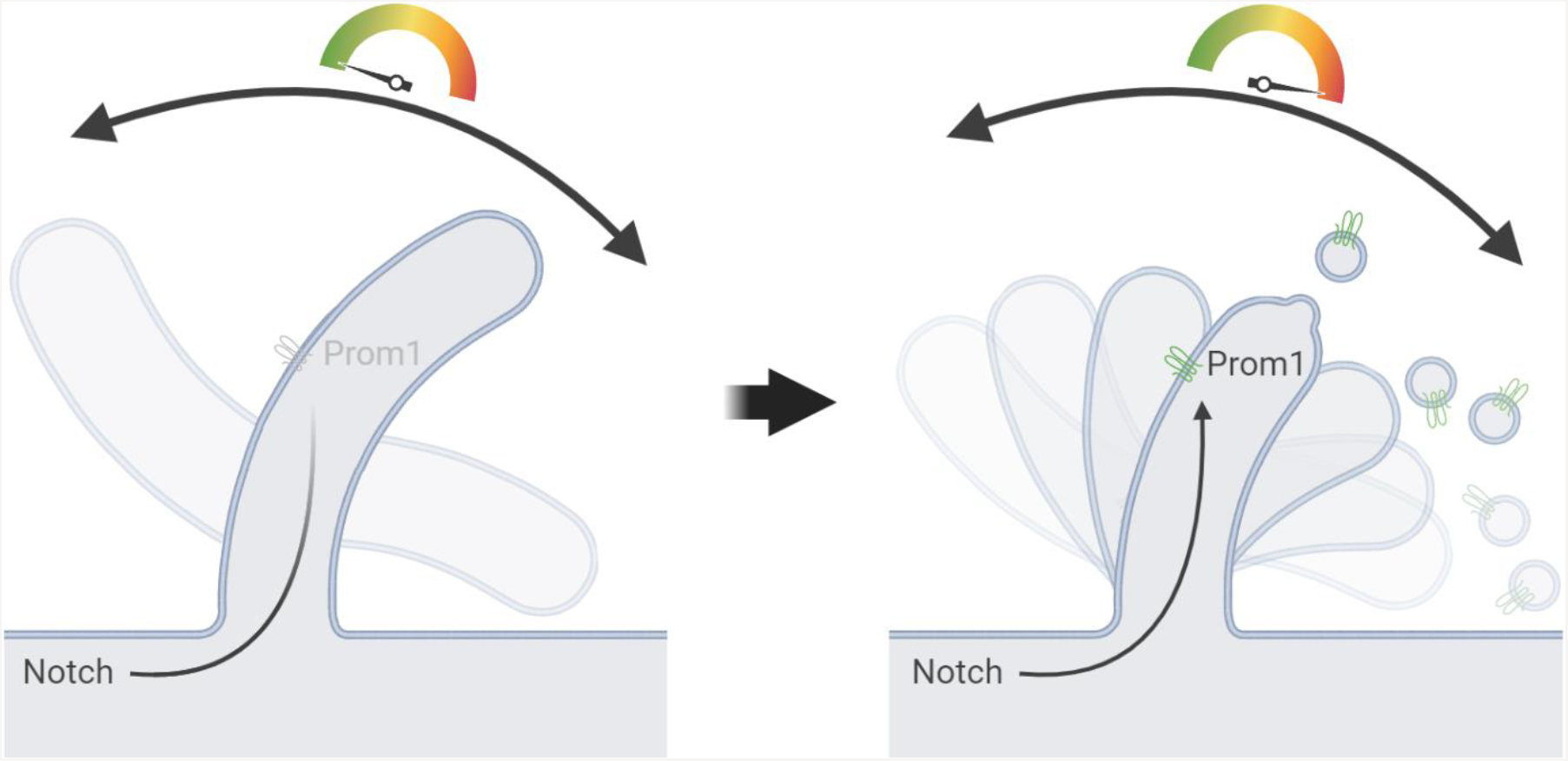
Proposed model for Prom1 regulation of the cilia phenotype in multiciliated cells. Low levels of Notch-signaling are present in cells committed to initiate the multiciliated fate. In these cells, Notch-signaling is involved in modulating the expression of cilia-related genes, such as Prom1, that contribute to the fine tuning of the ciliary phenotype. A variable underlying activation of Notch-signaling is responsible for the different levels of Prom1 expression along the airway, which modulate the ciliary phenotype by restricting cilia growth due to the increased shedding of Prom1-rich membrane particles from the multicilia. Through this mechanism, the expression of Prom1 in multiciliated cells contributes to the generation of diversity within this cell type. Illustration created with BioRender.com.

